# Metabolic reprogramming by endothelial ANGPTL4 depletion protects against diabetic kidney disease

**DOI:** 10.1101/2025.05.08.652142

**Authors:** Swayam Prakash Srivastava, Han Zhou, Rachel Shenoi, Myshal Morris, Begoña Lainez-Mas, Yuta Takagaki, Barani Kumar Rajendran, Ocean Setia, Binod Aryal, Keizo Kanasaki, Daisuke Koya, Ken Inoki, Alan Dardik, Carlos Fernández-Hernando, Gerald I. Shulman, Julie E. Goodwin

**Author notes:** Correspondence: Swayam Prakash Srivastava, & Julie E. Goodwin, Department of Pediatrics, Yale University School of Medicine, CT USA, 06520.

## Abstract

The role of cell-specific ANGPTL4 is not well known in the context of ECs, specifically in pathological angiogenesis and its relation to diabetic kidney disease. Here, we demonstrate that endothelial ANGPTL4 is required to induce a metabolic phenotype that favors mesenchymal activation in ECs and tubules in diabetic conditions. Diabetes accelerates mesenchymal activation and fibrogenesis in control mice however, the same effects were not observed in endothelial-cell specific knock out mice. This mesenchymal activation in diabetes is directly linked with pathological neovascularization, endothelial leakage, lipid and glycolysis metabolite load, de novo lipogenesis (DNL) and related mitochondrial damage, activation of the immune system, c-GAS-STING activation and transcription of pro-inflammatory cytokines. However, endothelial ANGPTL4-depleted mice had stable vessels, improved levels of lipid and glucose metabolism, suppressed levels of DNL, restored mitochondrial function, and mitigated levels of c-GAS-STING-mediated inflammation. Moreover, Inhibition of DNL, and STING via small molecule inhibitors suppressed pathological neovascularization and endothelial leakage, normalized fatty acid oxidation and reduced pathological glycolysis and de novo lipogenesis (DNL). These data demonstrate the crucial roles of endothelial ANGPTL4 in regulating pathogenic angiogenesis in the renal vasculature during diabetes.

## Introduction

Diabetic kidney disease (DKD) is the most common cause of end-stage renal disease^1–3^. Though there are several mechanisms reported to lead to renal fibrosis, the ultimate pathophysiology is still not clear and remains elusive^4–7^. Fibrosis is characterized by deposition of collagens, activation of myo-fibroblast formations, and activation of inflammatory cells and integrins in the interstitial space^7–14^. The origin and proliferation of renal myo-fibroblasts play a critical role in the development of diabetic kidney fibrosis^5–7,15,16^. Five major mechanisms have been proposed for the origin of renal myofibroblasts: 1) renal myofibroblasts can originate through epithelial cells by undergoing epithelial-to-mesenchymal transition (EMT) (though this idea is somewhat controversial and the term partial EMT has been proposed), which contributes less than 5% of total renal myofibroblasts^8,9,17–20^, 2) renal M2-type macrophages can transform into myofibroblasts through macrophage-to-mesenchymal transition^21,22^, 3) myofibroblasts can be derived from bone marrow cells^23^, 4) upon inflammation or stimuli, kidney resident fibroblast may activate and gain myofibroblast features^17^, 5) ECs can contribute to the generation of myofibroblasts through endothelial-to-mesenchymal transition (EndMT) processes^5,18,24–27^. EndMT-derived intermediate cell types which co-express both endothelial markers and myofibroblast markers are the most important cell type in the fibrotic process, since they have the ability to proliferate and influence neighboring cells’ physiology and homeostasis^5,8,26,28^. Alterations in the permeability and leakage of ECs and rupture of the connections between ECs and pericytes or disturbances in pericyte coverage are the most common cause of defective angiogenesis in tumors, however less is known about this process in diabetes^26,27,29^.

Biological pathways such as activated transforming growth factor-β (TGFβ) signaling^30^, Notch signaling^31–33^, Wnt signaling^34–36^, and Hedgehog signaling^37,38^ lead to disruption in central metabolism and cause mesenchymal metabolic shifts in diabetic kidneys, ultimately resulting in deposition of collagens, extracellular matrix proteins, fibronectin, vimentin, and N-cadherin in interstitial spaces.

Endothelial cell specialization in each organ is essential for executing tissue-specific functions^39–42^. Kidneys have a unique vasculature to regulate blood pressure, electrolyte homeostasis, pH balance and even blood cell production^42^. Most importantly, kidney ECs signal to other neighboring cell types, such as podocytes and tubules, to regulate their growth and proper function^42^. Any disturbances in the regulation between endothelial-podocyte crosstalk or endothelial-tubular cell crosstalk affect kidney health and function, however, endothelial-tubular cell crosstalk is one of the least studied areas in diabetes^43–45^. Renal angiogenesis is characterized by ECs losing their quiescence, supressing barrier function, proliferating, degrading extracellular matrix, altering their adhesive properties, migrating, forming tube-like structures and eventually developing into new blood vessels^46–50^. Renal angiogenesis is a complex process involving several molecular and cellular events which lead to specific programs of gene expression to support an angiogenic response which has recently been shown to be driven by a metabolic switch^49–54^. For instance, diabetes-associated hypoxia promotes vessel growth by upregulating many pro-angiogenic pathways and reinforcing endothelial cell (EC) glycolytic metabolism.

Angiopoietin-like 4 (ANGPTL4) is part of a family of eight proteins from related gene products that are structurally similar to the angiopoietins, is known to inactivate lipoprotein lipase (LPL) and is primarily expressed in adipose tissue, liver, and skeletal muscle^55–58^. ANGPTL4 participates in an array of biological functions, including the regulation of lipid and glucose metabolism, wound healing, hematopoietic stem cell expansion, angiogenesis, fibrosis and proteinuria^29,56,59,60^. Moreover, ANGPTL4 has been found to be associated with many diseases, including cancer, diabetes, atherosclerosis and renal fibrosis^59–62^. ANGPTL4 was discovered by screening of novel PPAR-γ targets for novel fasting-induced factors from the liver and PCR screening to identify novel angiopoietin-related proteins^63,64^. ANGPTL4 is also expressed in the vascular endothelium where it has been reported to regulate many *a priori* non-metabolic functions^65^. Indeed, given its similarity in structure with angiopoietins, ANGPTL4 attracted attention as a potential angiogenic and vascular permeability factor, especially given its control of a multitude of EC functions including vascular inflammation, nitric oxide production, angiogenesis, and vascular permeability^66–69^. Most of these studies only considered the paracrine actions of ANGPTL4 on ECs and neglected the potential of a specific lipase-dependent autocrine role of endothelial ANGPTL4. ANGPTL4 is a critical factor in determining plasma triglyceride (TG) levels in endothelial-bound LPL, which hydrolyzes triglycerides (TG), releases non-esterified fatty acids, and also promotes tissue uptake of non-esterified fatty acids^56,57,70^. Interestingly, EC expression of ANGPTL4 is induced by hypoxia, and was initially described to induce a strong pro-angiogenic response^65,71^. However, contradictory results have been reported regarding its contribution to angiogenesis and vascular integrity^67,69,72–74^. These apparent discrepancies seem to derive from the use of different experimental approaches (i.e., the use of in vitro systems with different forms of ANGPTL4), and the use of global knockout animal models to address the effect on the endothelium. Clement et al.^75^ observed two forms of ANGPTL4: a high-isoelectric pro-proteinuric form, found only in glomeruli which is hyposialylated, and a sialylated, neutral isoelectric form that is secreted from adipose tissue, liver, and skeletal muscle. Conversion of the hyposialylated form to the sialylated form by treatment with N-acetyl-d-mannosamine suppresses proteinuria and the disease phenotype in a mouse model of nephrotic syndrome^75^. A study using a podocyte-specific transgenic overexpression model has demonstrated higher podocyte-specific Angptl4 secretion with more than ∼500-fold increased proteinuria, loss of glomerular basement membrane (GBM) charge and foot process effacement, whereas transgenic overexpression in adipose tissue resulted in increased circulating Angptl4 and hypertriglyceridemia but no proteinuria, suggesting opposing effects of Angptl4 in renal tissues as compared to extrarenal tissues^75–77^. The CD1; *db/db* mouse, which is a mouse model of type 2 diabetes, mimics the renal fibrotic features of human DKD, has significantly higher expression levels of kidney *Angplt4*, suggesting a potential role of Angplt4 in DKD^78^. In addition, deficiency of ANGPTL4 has been shown to lower circulating lipids and reduce risk of coronary artery disease in humans^79,80^.

The results from our group demonstrated that tubule- and podocyte-depleted ANGPTL4 mice reprogram tubule and podocyte lipid metabolism and were protective in a mouse model of DKD^59^. Metabolic reprogramming, which is generally characterized by alterations in metabolic shifts in different cell types, is one the predominant features of mesenchymal activation in injured kidney cells ^35,81–85^. It is still not well known how these metabolic shifts are regulated or how they stimulate mesenchymal activation in ECs. However, it has been shown that suppression of fatty acid oxidation (FAO) in injured epithelial cells is related to development of fibrosis and that aberrant glycolysis in injured ECs contributes to mesenchymal activation processes^24,81,82,86^. To investigate this question further we used endothelial-specific ANGPTL4 knockout mice. We determined that EC-specific ANGPTL4 knockout favors lipid oxidation and suppresses glycolysis, which is associated with suppressed c-GAS-STING activation and increased expression of SIRT1 in endothelial cells. Overall we conclude that EC-ANGPTL4 depletion participates in metabolic reprogramming that augments FAO and is involved in renal EC angiogenic function and permeability.

## Material & Methods

### Reagents and antibodies

Rabbit polyclonal antibody against ANGPTL4 (#40-9800), rabbit polyclonal TFAM (Cat:PA5-29571), mouse monoclonal αSMA (Cat: MA1-26017) and rabbit anti-BIODIPY^TM^ FL (Cat: A5770) were purchased from Invitrogen. Mouse monoclonal anti-αSMA (Cat: A5228) was purchased from Sigma (St Louis, MO). Goat specific anti-CD31 (Cat: AF3628) antibody was purchased from R&D System. Rat specific anti-mouse CD31 (Cat: 550274) was purchased from BD Pharmingen. Anti-TGFβR1 (Cat: ab31013), PPARα (Cat: ab215270), rabbit polyclonal anti-TGFβRII (Cat:ab61213), mouse monoclonal anti-vimentin (RV202) (Cat:ab8978), rabbit polyclonal anti-αSMA (ACTA2) (Cat:ab5694) and rabbit polyclonal p-65 of NFkB (AB16502) antibodies were purchased from Abcam (Cambridge, UK). Carnitine palmitoyl transferase 1a (CPT1a) (Cat:12252), rabbit polyclonal anti-E-cadherin antibody (24E10) (Cat:3195) and rabbit polyclonal PKM2 (D78A4)-XPCR (cat:4053S) antibodies were purchased from Cell Signaling Technology (Danvers, MA, USA). Rabbit polyclonal PFKFB3 (Cat: 13763-1-AP) and rabbit polyclonal p-IKBα-Serine32/36 (Cat: 82349-1-RR) were purchased from Peprotech. Rabbit polyclonal anti-smad3 (#9513) antibody was obtained from Cell Signaling Technology (Danvers, MA, USA). Fluorescence-, Alexa fluor 647-, and rhodamine-conjugated secondary antibodies were obtained from Jackson ImmunoResearch (West Grove, PA).

### Animal Experimentation

All experiments were performed according to a protocol approved by the Institutional Animal Care and Use Committee at Yale University and Kanazawa Medical University Japan. The experiments at Yale University were in accordance with the National Institute of Health (NIH) Guidelines for the Care of Laboratory Animals and experiments at Kanazawa Medical University were carried out in accordance with Kanazawa Medical University animal protocols (#2014-89; #2013-114 and #2014-101) and approved by the Institutional Animal Care and Use Committee (IACUC). For the cell specific loss-of-function of Angptl4, we bred the Angptl4 flox/flox mice with Tie1-cre mice (emut) to generate mice with deletion of Angptl4 in ECs. All mice were on the C57BL/6 background. Induction of diabetes in CD-1 mice and C57BL/6 mice was performed according to previously established experimental protocols^81,87–91^. A single IP dose of 200 mg/kg STZ was used to induce diabetes in CD-1 mice. In mice on the C57BL/6 background, diabetes was induced in 10-week-old male mice with five consecutive intraperitoneal (IP) doses of streptozotocin (STZ) 50 mg/kg in 10 mmol/L citrate buffer (pH 4.5). For inhibitor studies, 16 week-STZ treated diabetic control and diabetic emut mice were randomly assigned to receive endothelial lipase inhibitor (XEN-445), FASN inhibitor (FASNALL-FASi) or STING inhibitor (STINGi). XEN-445 was provided to 16 week-STZ-treated diabetic control and diabetic emut mice using a dose of 10 mg/kg through oral gavage once a day for 8 weeks^29,92^. FASN inhibitor (FASi) was provided to 16 week-STZ-treated diabetic control and diabetic emut mice using a dose of 10 mg/kg i.p. thrice in a week for 8 weeks^93^. In other experiments, STING inhibitor-C-176 (STINGi) was injected intraperitoneally 7.5 ul (750nmol) per mouse or DMSO dissolved in 85μl corn oil thrice weekly for 8 weeks in 16 week-STZ-treated diabetic control and diabetic emut mice^94^.

In another set of experiments, diabetic male mice were randomized to one of three groups sixteen weeks after induction of diabetes: (i) untreated, (ii) 2-deoxyglucose 500 mg/kg via intraperitoneal injection twice per week^95^, (iii) PKM2 activator (TEPP-46), dose 30 mg/kg via oral gavage or vehicle for four weeks^96^. Mice were treated for 4 weeks and compared to untreated diabetic mice. All mice had free access to food and water during experiments. Blood was obtained by retro-orbital puncture. Blood glucose concentration was measured using glucose-strips. Urine albumin levels were assayed using a Mouse Albumin ELISA Kit (Exocell, Philadelphia, PA). Tissue and blood were harvested at the time of sacrifice. Some kidneys were minced and stored at -80°C for gene expression and protein analysis. Other kidneys were placed immediately in optimal cutting temperature (OCT) compound for frozen sections or 4% paraformaldehyde for histologic staining. Prior to sacrifice, mice were weighed and blood glucose levels were measured using gluco-strips. Urine samples for albumin and creatinine levels were collected by using metabolic cages.

### Mouse model of unilateral ureteral obstruction (UUO)

UUO surgery procedure was performed as previously described^18,97^. Briefly, mice were anesthetized with isoflurane (3%–5% for induction and 1%–3% for maintenance). Mice were shaved on the left side of the abdomen, a vertical incision was made through the skin with a scalpel, and the skin was retracted. A second incision was made through the peritoneum to expose the kidney. The left ureter was tied twice 15 mm below the renal pelvis with surgical silk, and the ureter was then severed between the 2 ligatures. Then, the kidney was placed gently back into its correct anatomical position and sterile saline was added to replenish fluid loss. The incisions were sutured and mice were individually caged. Buprenorphine was used as an analgesic. A first dose was administered 30 minutes before surgery and then every 12 hours for 72 hours, at a dose of 0.05 mg/kg subcutaneously. Mice were sacrificed and kidneys were harvested after perfusion with PBS at 11 days after UUO. Contralateral kidneys were used as a nonfibrotic control for all experiments using this model.

### Transmission electron microscopy

For electron microscopy studies, mice were anesthetized and perfused with 4% paraformaldehyde and kidneys were isolated. Kidney samples (1 mm^3^) were fixed with cacodylic acid buffer containing 1 M cacodylic acid and 25% glutaraldehyde in PBS. After Epon-embedding, an RMC/MTX ultramicrotome (Elexience) was used to cut the tissues into ultrathin sections (60 to 80 nm). Sections were mounted and imaged using a Nikon TE 2000U electron microscope on copper grids and stained with lead citrate and 8% uranyl acetate. A Jeol 1200 EX transmission electron microscope (Jeol LTD) equipped with a MegaView II high-resolution transmission electron microscopy camera was used to observe the copper grids. Electron microscopy was performed by the Cellular and Molecular Physiology Core at Yale University. ImageJ was used for quantitative analysis of electron micrographs. Foot processes were measured from at least 50 μm of GBM for each mouse. Podocyte foot processes were analyzed by ImageJ. Images were blinded by assigning integer numbers before evaluation by someone other than the scorer.

### Morphological Evaluation

Glomerular surface area was measured in 10 glomeruli per mouse using ImageJ software. We analyzed PAS-stained glomeruli from each mouse using a digital microscope screen. Masson’s trichrome-stained images were evaluated by ImageJ software, and the fibrotic areas were estimated^98^.

### Sirius red staining

Deparaffinized sections were incubated with picrosirius red solution for 1 hour at room temperature. The slides were washed twice with acetic acid solution for 30 seconds per wash. The slides were then dehydrated in absolute alcohol three times, cleared in xylene, and mounted with a synthetic resin. Sirius red staining was analyzed using ImageJ software, and fibrotic areas were quantified.

### Immunohistochemistry

Paraffin-embedded kidney sections (5 μm thick) were deparaffinized and rehydrated (2 min in xylene, four times; 1 min in 100% ethanol, twice; 1 min in 95% ethanol; 45 s in 70% ethanol; and 1 min in distilled water), and the antigen was retrieved in a 10 mM citrate buffer pH 6 at 98 °C for 60 min. To block endogenous peroxidase, all sections were incubated in 0.3% hydrogen peroxide for 10 min. Immunohistochemistry was performed using the Vectastain ABC Kit (Vector Laboratories, Burlingame, CA and Abcam ab64238). Rabbit polyclonal CPT1a (Abnova; H00001374-DO1P; 1:100) antibody was purchased from Abnova, USA. Rabbit polyclonal PKM2 (Cell Signalling Technology Cat# 4053, RRID:AB_1904096; 1:100) was purchased from Cell Signalling. In negative controls, the primary antibody was omitted and replaced with blocking solution.

### Immunofluorescence

Paraffin-embedded kidney sections (5 μm thick) were used for immunofluorescence; double positive CD31/FITC-dextran, CD31/Vimentin, CD31/Fibronectin, CD31/PFKFB3, CD31/PKM2, CD31/CPT1a, CD31/αSMA, CD31/TFAM, CD31/DPP-4, CD31/β1-integrin, CD31/p-IKBα and CD31/p65-NFkB, E-cadherin/αSMA and E-cadherin/Vimentin. Paraffin-embedded kidney sections were deparaffinized and rehydrated (2 min in xylene, four times; 1 min in 100% ethanol, twice; 1 min in 95% ethanol; 45 s in 70% ethanol; and 1 min in distilled water), and the antigen was retrieved in a 10 mM citrate buffer pH 6 at 98 °C for 60 min. To block endogenous peroxidase, all sections were incubated in 0.3% hydrogen peroxide for 10 min and then blocked in 2% bovine serum albumin/PBS for 30 min at room temperature. Then, sections were incubated in primary antibody (1:100) for 16 h at 4°C and on the second day were washed in PBS (5 min) three times. Next, the sections were incubated with secondary antibodies (1:200 dilution) for 60 min, washed with PBS three times (10 min each), and mounted with mounting medium with DAPI (Vector Laboratories, Burlingame, CA). The stained sections were analyzed by fluorescence microscopy. For each mouse, original magnification of 400X was obtained from six different areas and quantified.

Frozen kidney sections (5 μm) were also used for immunofluorescence; double positive labelling with CD31/αSMA and E-cadherin/αSMA was measured. Briefly, frozen sections were dried and placed in acetone: methanol (1:1) solution for 10 min at − 30 °C. Once the sections were dried, they were washed twice in phosphate-buffered saline (PBS) for 5 min and then blocked in 2% bovine serum albumin/PBS for 30 min at room temperature. Then, sections were incubated in primary antibody (1:100) for 1 h and washed in PBS (5 min) three times. Next, the sections were incubated with secondary antibodies for 60 min, washed with PBS three times (10 min each), and mounted with mounting medium with DAPI (Vector Laboratories, Burlingame, CA). The stained sections were analyzed by fluorescence microscopy. For each mouse, original magnification of 400X was obtained from six different areas and quantified.

### Multiplex staining

Multiplex staining was analyzed according to the manufacturer’s instructions by using an Opal in situ kit (Waltham, MA, USA). Deparaffinized sections were labeled with E-cadherin (Opal 520 [TSA-FITC]) and Vimentin (Opal 670 [TSA-Cy5]) antibodies for EMT transition analysis while α-SMA (opal 520 [TSA-FITC]) and CD31 (opal 670 [TSA-Cy5]) antibodies were used for EndMT analysis. The cell nuclei were labeled with DAPI. In negative controls, the primary antibody was omitted and replaced with blocking solution.

### Proximity ligation Duolink in situ assay

Duolink In Situ kits were used to detect the proximity of DPP-4/integrin β1 and TGF-βR1/R2, as previously described^99,100^. Briefly, cells were passaged into 8-well culture slides (BD Biosciences) in growth medium. The cells were washed with PBS, fixed with 4% paraformaldehyde, and permeabilized with 0.2% Triton-X100. Then, blocking solution was used for 30 min at 37 °C, after which the cells were incubated in primary antibody (goat anti-integrin β1 (1:100) and rabbit anti-Angptl4 (1:100), or rabbit anti-TGFβR1 (1:500) and goat anti-TGFβR2 (1:500), or goat polyclonal DPP-4 (1:100) and rabbit anti-integrin β1 (1:100)) at 4 °C overnight. Cells were treated with two PLA probes for 1 h at 37°C, Ligation-ligase solution was added for 30 min at 37 °C, and amplification-polymerase solution was added for 100 min at 37 °C. The slides were then mounted with DAPI and analyzed by fluorescence microscopy. For each slide, original magnification of 400X was obtained from 6 different areas and quantified.

### Vascular Permeability assay

To determine vascular permeability, mice were intravenously injected with 100μl FITC-Dextran (70 kDa or 4 kDa, 25 mg/mouse)^29^. Extracellular vascular FITC-Dextran fluorescence was analyzed by a fluorescence microscope.

### Enzyme assays

Enzyme activity was analyzed in isolated ECs following the instructions from the standard kits (Biovision).

### Oil Red O Staining

Briefly, 8 µm kidney sections were dried at room temperature for 15 min and fixed in pre-chilled acetone solution for 10min. Slides were washed three times with PBS for 5 min and finally rinsed in 60% isopropanol for 5 min. Lipids were stained by incubating slides in fresh Oil Red O working solution for 60 min at room temperature and rinsed in 60% isopropanol for 5 seconds. Slides were washed three times with PBS and 5 min with water and counterstained with hematoxylin. Finally, slides were washed in 70% ethanol and mounted with mounting media, and immediately imaged.

### EndMT and EMT detection

Frozen sections (5 µm) were used for the detection of EndMT and EMT. Cells undergoing EndMT were detected by double-positive labeling for CD31 and αSMA and with vimentin. Cells undergoing EMT were detected by double-positive labeling for E-cadherin and αSMA and with vimentin. Sections were analyzed and quantified by fluorescence microscopy.

### Isolation of ECs

ECs from the kidneys of non-diabetic and diabetic mice were isolated using CD31 magnetic beads. Briefly, kidneys were isolated and minced into small pieces. Using a series of enzymatic reactions by treating the tissue with trypsin and Collagenase type I solution, a single cell suspension was created. The pellet was dissolved with CD31 magnetic beads, and the CD31-labeled cells were separated with a magnetic separator. The cells were further purified on a column. Cell number was counted by hemocytometer and cells were plated on 0.1% gelatin coated Petri dishes. Cell purity was measured by flow cytometry (BD FACSDiva) using PE-conjugated CD31 (BDB553373) and FITC-conjugated CD45 (BDB553079), both from Becton Dickinson (USA)^35^.

### Isolation of kidney tubular epithelial cells (TECs)

After sacrifice, kidneys from diabetic Angptl4^-/-^ and control littermates were excised and perfused with 4% paraformaldehyde (10 mL) followed by collagenase type II digestion (2 mg/mL). After digestion, the cortical regions of the kidneys were used for further processing. The cortical region was minced and digested in collagenase buffer for an additional 5 minutes at 37°C with rotation to release cells. Digested tissue and cell suspension were passed through a 70-μm cell strainer, centrifuged at 50 g for 5 min, and washed in PBS for 2 rounds to collect TECs. Isolated TECs were seeded onto collagen-coated Petri dishes and cultured in renal epithelial cell medium (C-26130, PromoCell) supplemented with growth factors for TEC growth^35^.

### Seahorse Flux Analysis of Kidneys

Basal oxygen consumption rates in kidneys from non-diabetic and diabetic wild-type and Angptl4^-/-^ mice were characterized using the Seahorse Flux Analyzer (Agilent) according to the manufacturer’s instructions. Briefly, mice were fasted overnight, sacrificed and perfused with 1x PBS. Whole kidneys were rapidly dissected, washed in 1x KHB buffer and cut into ∼2 mg pieces. Following dissection, kidney pieces were snapped into the wells of a XF Islet Capture Microplate containing 500 ml XF24 Assay Media (DMEM base media containing 1 mM pyruvate, 2 mM glutamine, 5.5 mM glucose and 100 mM palmitate, pH 7.4). Kidneys were equilibrated at 37°C for 1 h in a non-CO_2_ incubator and then assayed on a Seahorse XFe24 Analyzer (Agilent) following a 12-min equilibration period. Respiration rates were measured three times using an instrument protocol of 3-minute mix, 2-minute wait, and 3-minute measure. Flux rates were normalized to tissue weight. Experiments were repeated in 3 mice per condition.

### Triglyceride measurement

Renal triglycerides were measured using Triglyceride Calorimetric Assay kit (#10010303, Cayman). Approximately 50mg of kidney tissue was homogenized in NP40-substitute assay buffer containing protease and phosphatase inhibitors. Homogenates were collected after centrifuging at 10,000g for 10min at 4°C.

### Cholesterol measurement

Renal cholesterol was quantitatively measured using established methods^93^. Approximately 20 mg of kidney tissue was homogenized in 300 µl of chloroform:isopropanol:NP-40 (7:11:0.1) in a microcentrifuge. The homogenate was centrifuged at 15000g for 10min, and the liquid layer was collected into a new microtube. The supernatant was air dried at 50 C to remove chloroform and the samples kept under vacuum pressure for 30 min to remove trace organic solvent. The dried lipids were dissolved in 200 µl of assay buffer and cholesterol measurements were performed as per the manufacturer’s protocol for the kit.

### ATP measurement

ATP content was determined using the ATP Colorimetric Assay kit (Biovision), following the manufacturer’s instructions.

### mRNA isolation and qPCR

Total RNA was isolated using standard Trizol protocol. RNA was reverse transcribed using the iScript cDNA Synthesis kit (Bio-Rad) and qPCR was performed on a Bio-Rad C1000 Touch thermal cycler using the resultant cDNA, as well as qPCR Master mix and gene specific primers. The list of mouse primers used is in **Table S1**. Results were quantified using the delta–delta-cycle threshold (Ct) method (ΔΔCt). All experiments were performed in triplicate and 18S was utilized as an internal control.

### *In vitro* experiments and transfection

Human HK-2 cells were used at passages 4-8 and cultured in epithelial basal media with growth factors and 10% serum. Human Angptl4 siRNA (Invitrogen) was used at a concentration of 40 nM for 48 h to effectively knock down Angptl4. Cells were treated with or without TGFβ2 (10 ng/ml) for 48h. In a second set of experiments, isolated ECs from diabetic control and diabetic emut mice were cultured. The conditioned media (CM) from diabetic control and diabetic emut were subsequently transferred to cultured isolated tubules. In another experiment, Human HK-2 cells were cultured in DMEM and Keratinocyte-SFM (1X) (Life Technologies Green Island NY) media, respectively. When the cells on the adhesion reagent reached 70% confluence, 10 ng/ml recombinant human TGFβ1 was placed in the serum-diluted medium for 48 h.

### Fatty acid uptake

Cultured isolated kidney ECs were incubated with medium containing 2 μCi [^14^C]- palmitate. [^14^C]-palmitate uptake was measured by liquid scintillation counting.

### Fatty Acid Oxidation

Cultured isolated kidney ECs were incubated with medium containing 0.75 mmol/L palmitate (conjugated to 2% free fatty acid–free BSA/[^14^C] palmitate at 2 μCi/mL) for 2 h. One mL of the culture medium was transferred to a sealable tube, the cap of which housed a Whatman filter paper disc. ^14^CO_2_ trapped in the media was then released by acidification of media using 60% perchloric acid. Radioactivity that had become adsorbed onto the filter discs was quantified by liquid scintillation counting.

### Statistical analysis

All values are expressed as mean ± SEM and analyzed using GraphPad Prism 7 (GraphPad Software, Inc., La Jolla, CA). One-way ANOVA followed by Tukey’s test and two-way ANOVA were employed to analyse significance when comparing multiple independent groups. The post hoc tests were run only if F achieved P< 0.05 and there was no significant variance in homogeneity. In each experiment, N represents the number of separate experiments (in vitro) or the number of mice (in vivo). Technical replicates were used to ensure the reliability of single values. Data analyses were blinded. The data were considered statistically significant at p < 0.05.

## Results

### Endothelial dysfunction in diabetes is related to higher ANGPTL4 gene expression

The streptozotocin-induced diabetic CD-1 mouse model is the established mouse model for studying DKD^81,89,90^. 24-week post streptozotocin injected CD-1 mice show elevated levels of collagen deposition and fibrosis in kidneys **(Figure 1A).** Immunofluorescence studies reveal that this increased renal fibrosis is associated with higher levels of α-SMA in CD-1 positive cells, suggesting increased EndMT in these diabetic kidneys **(Figure 1B).** Endothelial density, as represented by increased density of CD31-positive cells was increased in diabetic kidneys **(Figure 1C).** Endothelial PFKFB3 and PKM2 were analyzed by measuring co-labeled PFKFB3 and PKM2 in CD31 positive cells in control and diabetic kidneys **(Figure 1C).** Diabetic kidneys had higher levels of PFKFB3 and PKM2 compared to those in nondiabetic controls **(Figure 1C).** To study EC permeability in diabetes, we injected control and diabetic mice with FITC-dextran and then isolated the kidneys for analysis. FITC-dextran levels in CD31 positive cells were higher in diabetic kidneys, indicating greater permeability **(Figure 1D).** Endothelial CPT1a was analyzed by measuring co-labeled CPT1a in CD31 positive cells in control and diabetic kidneys **(Figure 1D).** Diabetic kidneys had lower levels of CPT1a expression compared to those of nondiabetic controls **(Figure 1D).** Isolated ECs from control and diabetic kidneys were subjected to a TGFβR1-TGFβR2 interaction assay using the duo-link proximity ligation method, and we found increased TGFβR1 and TGFβR2 interactions in diabetic ECs when compared to control ECs **(Figure 1E).** Relative gene expression analysis of FAO-related genes such as *CPT1a* and *PPARα* were downregulated whereas expression of fatty acid transporter proteins *FATP1, FATP4, and CD36* was upregulated in diabetic ECs. The expression of mesenchymal markers including vimentin, *FSP-1* and glucose transporter protein 1 (*GLUT1*) was significantly upregulated in diabetic ECs **(Figure 1F).** Lactate levels and glycolysis intermediates such as pyruvate, glucose-6-PO_4_, fructose-6-PO_4_, phosphoenolpyruvate and key glycolysis enzymes such as hexokinase, phosphofructokinase, and pyruvate kinase were higher in diabetic ECs while pyruvate dehydrogenase (PDH) enzyme activity was lower in diabetic ECs **(Figure 1G-I).** Glucose uptake and radiolabeled palmitate uptake (lipid uptake) were higher, while palmitate oxidation was suppressed, in diabetic ECs when compared to control cells **(Figure 1J-K).** Relative gene expression analysis of the lipoprotein lipase regulator (LPL) *ANGPTL4* was significantly upregulated, while *ANGPTL2* and *ANGPTL7* gene expression were less elevated, in diabetic ECs when compared to control ECs **(Figure 1L).** Taken together, these data suggest defective endothelial metabolism and function is present in diabetes and is associated with upregulated *ANGPTL4* gene expression.

**Figure 1:**
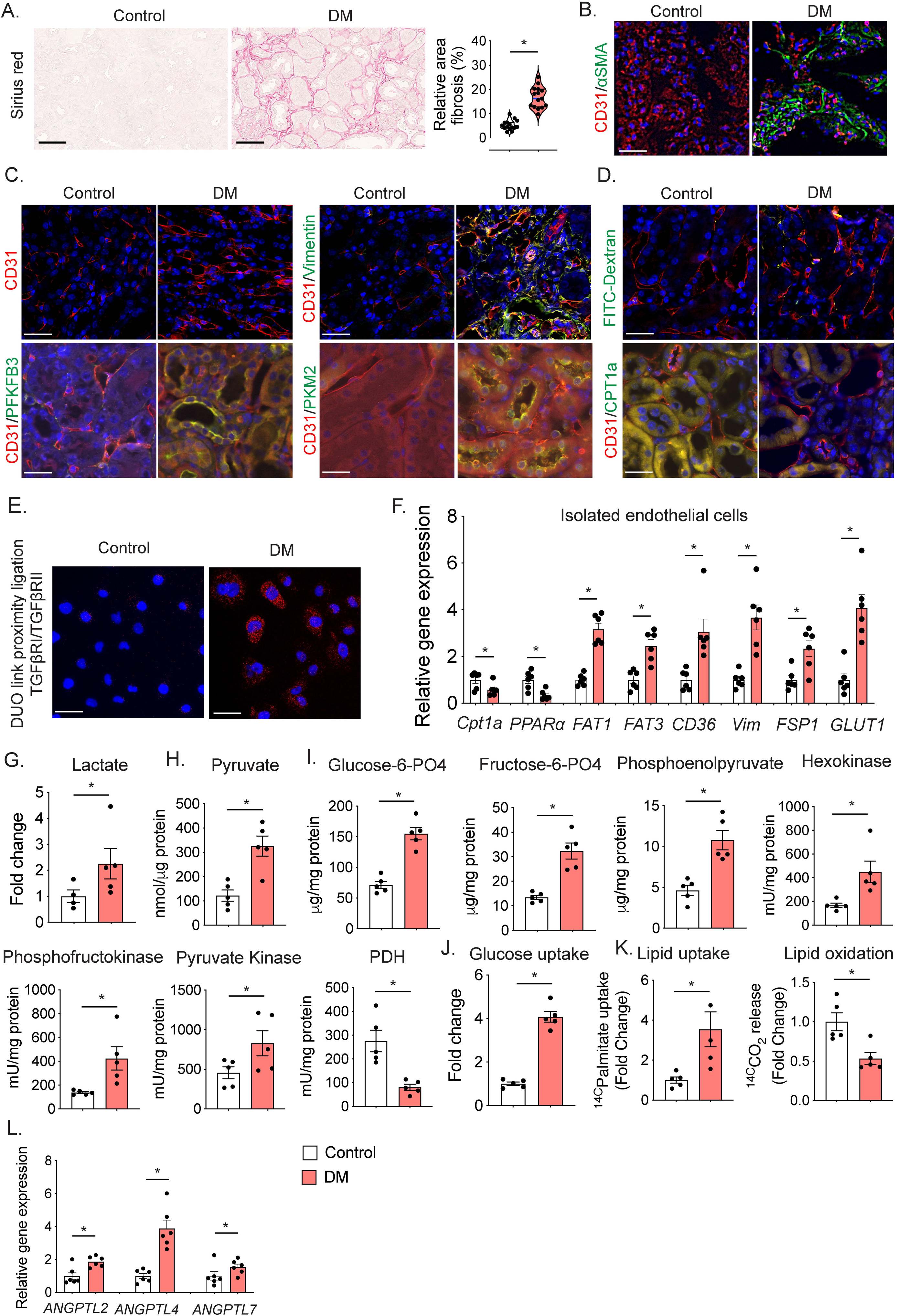
Mesenchymal and fibrogenic transformation is associated with defective lipid and glucose metabolism and related endothelium dysfunction in diabetes. **A.** Sirius red staining in kidneys of 28-week-old control and streptozotocin-induced diabetic CD-1 mice. Relative area of fibrosis was calculated using ImageJ. Representative images are shown. N=15/group. Scale bar 100 μm. Original magnification 40x. **B.** CD31/αSMA co-labelling was analyzed in kidneys of 28-week-old control and streptozotocin-induced diabetic CD-1 mice using immunofluorescence. Representative microphotographs are shown. FITC-αSMA , Rhodamine-CD31 and DAPI-blue. Original magnification 40x. N=8/group. Scale bar 50 μm. **C.** Endothelial density and mesenchymal activation in ECs were analyzed by CD31 staining and CD31/vimentin co-staining by immunofluorescence in the kidneys of 28-week-old control and streptozotocin-induced diabetic CD-1 mice. CD31/PFKFB3, CD31/PKM2 co-immunolabeling was also analyzed. FITC-vimentin, PFKFB3 and PKM3; Rhodamine-CD31 and DAPI-blue. Original magnification 40x. N=8/group. Scale bar 50 μm. Representative microphotographs are shown. **D.** Endothelial permeability was measured by analyzing FITC-dextran by using immunofluorescence. CD31/CPT1a was also analyzed. FITC-FITC-dextran, CPT1a; Rhodamine-CD31 and DAPI-blue. Original magnification 40x. N=6/group. Representative images are shown. Scale bar 50 μm. **E.** TGFβRI and TGFβRII proximity was analyzed by Duo-link proximity ligation in kidney sections from control and diabetic mice. Representative images are shown. N=6/group. Scale bar 50 μm. **F.** Relative mRNA expression levels of *CPT1a*, *PPARα*, , *FAT1*, *FAT 3*, and *CD36*, *Vimentin*, *FSP1* and *GLUT1* were measured in isolated ECs from nondiabetic and diabetic control and emut mice kidneys using quantitative PCR. 18S was used as an internal control to normalize the data. **G.** Lactate measurement, **H.** Pyruvate measurement, **I.** Glucose-6-PO_4_, fructose-6-PO_4_, phosphoenolpyruvate, hexokinase activity, phosphofructokinase activity, pyruvate kinase activity and pyruvate dehydrogenase enzyme activity measured in isolated ECs from control and diabetic kidneys, using available kits following their instructions. N=4∼5/group. **J.** Glucose uptake was measured in isolated ECs using the calorimetric method. N=5/group. **K.** Lipid uptake and lipid oxidation were measured as ^14C^palmitate uptake and ^14C^palmitate oxidation using established methods. CPM was counted using a scintillation counter. **L.** Relative mRNA expression levels of *ANGPTL2*, *ANGPTL4*, and *ANGPTL7* were measured in isolated ECs from diabetic and control kidneys and analyzed by qPCR. 18S was used as an internal control. Data are presented as mean±SEM. Student-t-test was used to calculated statistical significance between the groups.

### Metabolic reprogramming by endothelial ANGPTL4 depletion protects against DKD

Mice with conditional alleles (Angptl4^fl/fl^) were generated by removal of the gene trap cassette with Flp recombinase using the original tm1a mice^59,101^. These mice have a cassette containing the mouse En2 splicing acceptor, LacZ, and promoter-driven neomycin resistance gene and were inserted in the *Angptl4* gene. The initial allele (tm1a) generates a null allele through splicing to the LacZ trapping element. To test the role of *Angptl4* in ECs, we generated endothelial-specific Angptl4 mutant mice (emut) by crossing Angptl4^fl/fl^ mice with mice carrying the *Tie1 Cre* driver (Angptl4^fl/fl^; *Tie1 Cre+*) **(Figure 2A)**. In Angptl4^fl/fl^ mice, the expression of the LacZ reporter gene is regulated by the ANGPTL4 promoter gene in the original tm1a mice, and expression of ANGPTL4 can be detected using an antibody against β-galactosidase (β-gal). Therefore, we analyzed β-gal staining in the emut mice. Emut mice demonstrated higher expression of β-gal-CD31 co-labeled cells compared to littermate controls **(Figure S1A)**. Emut mice had significantly suppressed *Angptl4* expression in isolated ECs when compared to control mice **(Figure S1B)**. Diabetes was induced in both genotypes and mice had free access to food and water for 20 weeks **(Figure 2A).** Nondiabetic and diabetic emut mice did not show any remarkable alterations in body weight or blood glucose compared to nondiabetic control and emut mice respectively, whereas kidney weight and albumin-to-creatinine ratios (ACR) were remarkably suppressed in diabetic emut mice compared to the diabetic control mice **(Figure 2B-C)**. Moreover, the diabetic emut mice had reduced tubular damage, area of fibrosis, collagen accumulation and glomerulosclerosis compared to the diabetic control mice, as observed by kidney histology **(Figure 2C)**. Glomerular ultrastructure was analyzed by transmission electron microscopy and diabetic control mice displayed some podocyte foot process effacement while diabetic emut mice demonstrated this to a lesser extent **(Figure 2D)**. The phenotype of the emut mice was also evaluated using an accelerated model of nephropathy, which combines the effects of both the streptozotocin-induced diabetic model as well as the UUO model^97^. Mice were subjected to UUO followed by 4 daily doses of streptozotocin and euthanasia on day 11. There was less fibrosis in the kidneys of UUO-operated emut mice compared to those from UUO-operated control mice **(Figure S2)**.

**Figure 2:**
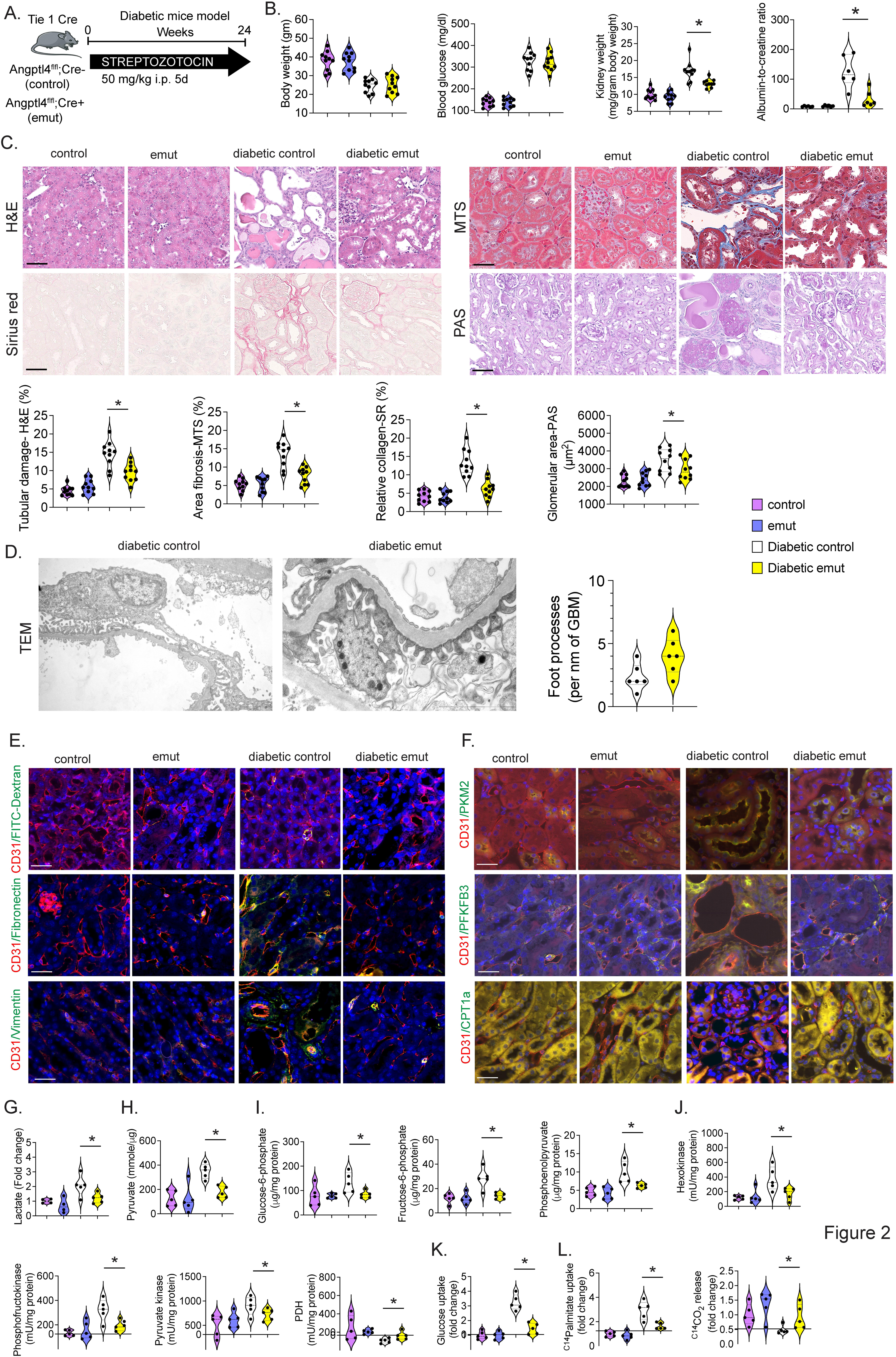
Endothelial ANGPTL4 depletion protects against DKD. **A.** Schematic chart showing a protocol for induction of diabetes. **B.** Body weight, blood glucose, kidney weight/body weight and urine albumin-to-creatine ratio. Blood glucose was analyzed using glucose strips. N=10/group. **C.** Kidney histology including H&E, Masson trichome staining (MTS), Sirius red (SR) and Periodic acid-schiff staining (PAS) were performed in kidneys. Representative images are shown. Tubular damage was analyzed from H&E, relative area fibrosis was analyzed from MTS, relative collagen was analyzed from SR and glomerular surface area was measured from PAS images. Original magnification 40x. N=10/group. Scale bar 50 μm. **D.** Representative transmission electron microscopy images. N= 3/group. Relative density of podocyte foot processes was calculated using ImageJ. **E.** Endothelial permeability was measured by analyzing FITC-dextran by immunofluorescence. FITC-FITC-dextran, Rhodamine-CD31 and DAPI-blue. Original magnification 40x. N=6/group. Representative images are shown. Scale bar 50 μm. **F.** Immuno-co-labelling of CD31/Vimentin, CD31/Fibronectin, CD31/PFKFB3, CD31/PKM2 and CD31/CPT1a was performed. FITC-Vimentin, Fibronectin, PFKFB3, PKM2, CPT1a; Rhodamine-CD31; DAPI-blue. Representative images are shown. N=6/group. Original magnification 40x. Scale bar 50 μm. **G.** Lactate measurement, **H.** Pyruvate measurement, **I.** Glucose-6-PO_4_, fructose-6-PO_4_, phosphoenolpyruvate, **J.** Hexokinase activity, phosphofructokinase activity, pyruvate kinase activity and pyruvate dehydrogenase enzyme activity measured in isolated ECs from control and diabetic kidneys. N=4-5/group. **K.** Glucose uptake assay in isolated ECs using the calorimetric method. N=5/group. **L.** Lipid uptake and lipid oxidation were measured as ^14C^palmitate uptake and ^14C^palmitate oxidation using established methods. CPM was counted using a scintillation counter. Data are represented as fold change. Data are presented as mean±SEM. One-way-Anova Tukey’s test was used to calculated statistical significance among the groups.

To analyze EC permeability we injected FITC-dextran via tail vein in nondiabetic and diabetic control and emut mice. Nondiabetic mice did not demonstrate any phenotype while the kidneys from diabetic control mice had higher FITC-dextran staining in CD31-positive cells, which was partially suppressed in the kidneys from diabetic emut mice **(Figure 2E)**. The kidneys of diabetic mice had higher increased EndMT and fibrosis as evidenced by higher vimentin and fibronectin staining in CD31-positive cells while the kidneys of diabetic emut mice demonstrated suppression of these processes **(Figure 2E,F)**. EndMT and fibrosis in the kidneys of diabetic controls were associated with higher levels of glycolysis markers such as PFKFB3 and PKM2 and suppressed expression of the fatty acid carrier protein CPT1a while kidneys from diabetic emut mice demonstrated the opposite result **(Figure 2F)**. Lactate levels and glycolysis intermediates such as pyruvate, glucose-6-PO_4_, fructose-6-PO_4_, phosphoenolpyruvate and key glycolysis enzymes such as hexokinase, phosphofructokinase, and pyruvate kinase were higher while pyruvate dehydrogenase (PDH) enzyme activity was lower in isolated ECs from diabetic controls while the reverse pattern was observed in isolated ECs from diabetic emut mice **(Figure 2G-J).** Glucose and lipid uptake were higher and palmitate oxidation was suppressed in ECs from diabetic controls while again the opposite effects were found in isolated ECs from diabetic emut mice **(Figure 2K-L).** Since ANGPTL4 inactivates local lipase and endothelial lipase (EL) in ECs, we analyzed the effect of endothelial lipase inhibition in diabetic control and emut mice. For this purpose we utilized the EL-specific inhibitor XEN-445 and treated diabetic mice for 3-weeks. We did not observe any remarkable change in body weight, blood glucose, kidney weight, ACR, tubular damage, area of fibrosis, collagen levels or glomerulosclerosis between the two genotypes (**Figure S3**).

### Endothelial ANGPTL4-depleted mice suppress de novo lipogenesis in diabetic endothelium

Kidneys of diabetic control mice had significantly elevated triglycerides compared to those of nondiabetic mice while the level in kidneys of diabetic emut mice was suppressed **(Figure S4A).** There were no changes in the cholesterol levels between diabetic groups **(Figure S4B).** To study the role of de novo fatty acid synthesis and mesenchymal activation in diabetic ECs, we examined the expression of key genes in this pathway. Isolated diabetic ECs from control mice had up-regulated gene expression of *FASN, ACC, perilipin, SREBF1 and SCAP*. The fatty acid transporter proteins *FAT1, FAT4* and *CD36* were also upregulated in diabetic control ECs. The expression of all of these genes were comparatively suppressed in ECs isolated from diabetic emut mice, similar to non-diabetic endothelial cell levels **(Figure S4C).** Additionally, diabetic kidneys of both genotypes had elevated FASN in CD31-positive cells compared to nondiabetic kidneys, and kidneys of diabetic emut mice demonstrated suppression compared to kidneys of diabetic controls suggesting a key role of ANGPTL4 in the regulation of FASN in diabetic ECs **(Figure S4D)**. To investigate the role of FASN in EndMT, we tested an inhibitor specific for FASN (FASi) in diabetic mice of both genotypes **(Figure 3A)**. We treated with FASi for 8 weeks and analyzed physiological parameters and kidney histology **(Figure 3A)**. We did not observe any differences in the body weight or blood glucose between diabetic groups, however we found significant suppression of kidney weight and ACR in FASi-treated control mice compared to vehicle-treated control mice **(Figure 3B)**; there were no differences between FASi-treated diabetic emut mice and vehicle-treated diabetic emut mice **(Figure 3B)**. We found a significant reduction in the tubular damage, area of fibrosis, collagen levels and glomeruloscelrosis in FASi-treated diabetic control mice, but again no differences between diabetic emut mice**(Figure 3C)**. The phenotype of the FASi-treated mice was also evaluated using the accelerated UUO +streptozotocin fibrosis model^97^. In this model, FASi was provided to the UUO-operated control and emut mice which were also subjected to 4 daily doses of streptozotocin followed by euthanasia on day 11. FASi-treated control and emut mice had lessened fibrosis compared to UUO-operated control mice and emut mice **(Figure S5)**.

**Figure 3:**
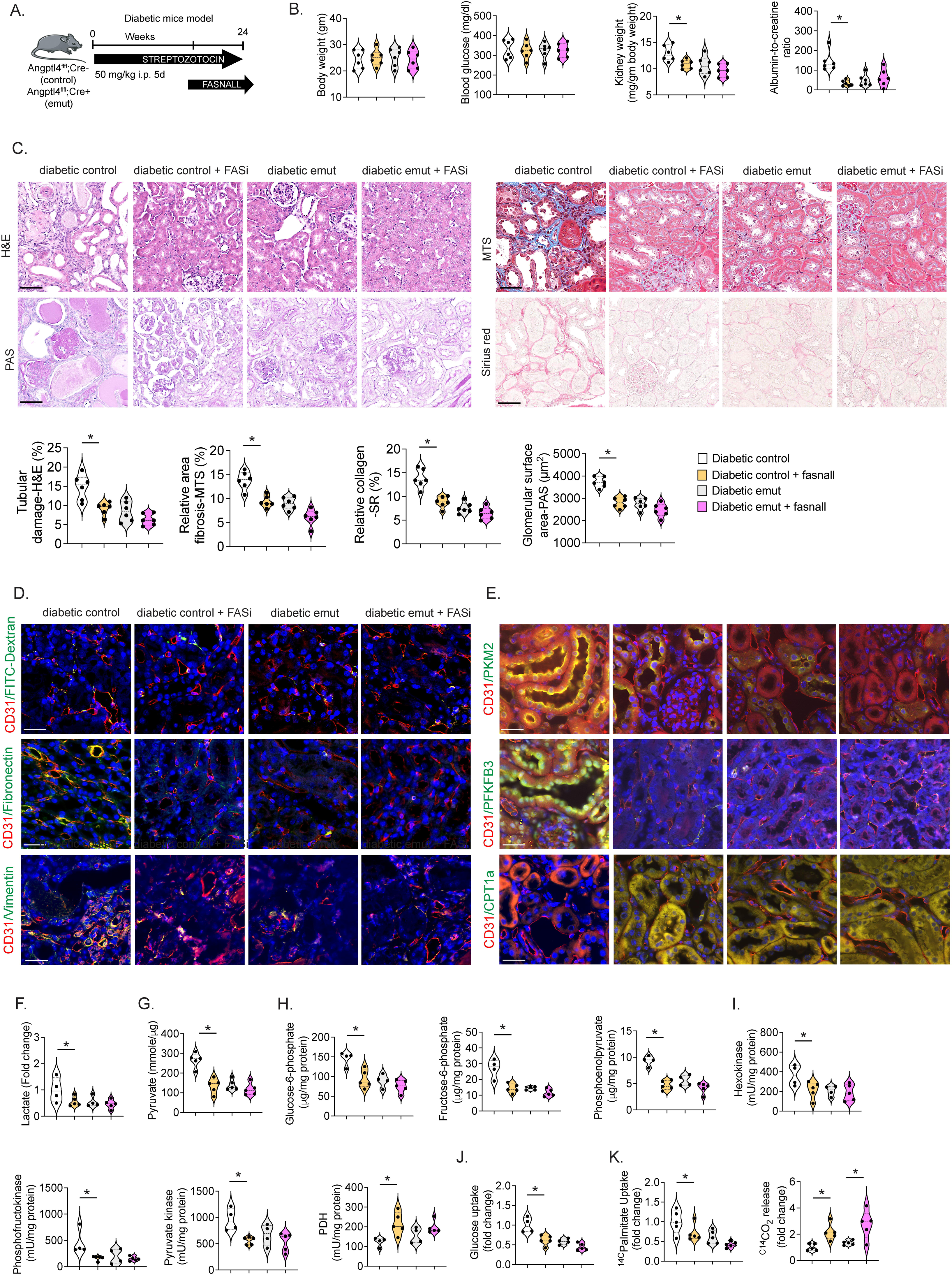
Fatty acid synthetase inhibition protects against DKD in diabetic control mice but not in diabetic emut mice. **A.** Schematic chart showing a protocol for the treatment of FASNALL (FASi) **B.** Body weight, blood glucose, kidney weight/body weight and albumin-to-creatinine ratio. N=6/group. **C.** Kidney histology in kidneys of FASi treated diabetic control and emut mice. Representative images are shown. Tubular damage was analyzed from H&E, relative area of fibrosis was analyzed from MTS, relative collagen was analyzed from SR and glomerular surface area was measured from PAS images. Original magnification 40x. N=6/group. **D.** Endothelial permeability was measured by analyzing FITC-dextran by immunofluorescence. FITC-FITC-dextran, Rhodamine-CD31 and DAPI-blue. Original magnification 40x. N=6/group. Representative images are shown. Scale bar 50 μm. **E.** Immuno-co-labelling of CD31/Vimentin, CD31/Fibronectin, CD31/PFKFB3, CD31/PKM2, CD31/CPT1a. FITC-Vimentin, Fibronectin, PFKFB3, PKM2, CPT1a, Rhodamine-CD31, DAPI-blue. Representative images are shown. N=6/group. Original magnification 40x. Scale bar 50 μm. **F.** Lactate measurement, **G.** Pyruvate measurement, **H.** Glucose-6-PO_4_, fructose-6-PO_4_, phosphoenolpyruvate, **I.** Hexokinase activity, phosphofructokinase activity, pyruvate kinase activity and pyruvate dehydrogenase enzyme activity measured in isolated ECs from the kidneys of FASi-treated diabetic control and emut mice. N=4-5/group **J.** Glucose uptake assay using the calorimetric method. N=4-5/group **K.** Lipid uptake and lipid oxidation were measured as ^14C^palmitate uptake and ^14C^palmitate oxidation using established methods. CPM was counted using a scintillation counter. Data is represented as fold change Data are presented as mean±SEM. One-way-Anova Tukey’s test was used to calculated statistical significance among the groups.

Endothelial cell leakage was suppressed in the FASi-treated diabetic control mice but no change was observed in the FASi-treated emut mice compared to vehicle-treated diabetic control or diabetic emut mice **(Figure 3D)**. The kidneys of FASi-treated diabetic mice had suppressed EndMT and fibrosis as evidenced by lower levels of vimentin and fibronectin staining in CD31-positive cells **(Figure 3E)**. This EndMT suppression was associated with reduced expression of PFKFB3 and PKM2 but increased expression of CPT1a compared to CD31-positive cells in vehicle-treated diabetic mice **(Figure 3E)**. Lactate, pyruvate, glucose-6-PO_4_, fructose-6-PO_4_, phosphoenolpyruvate and key glycolysis enzymes such as hexokinase, phosphofructokinase and pyruvate kinase were lower while PDH enzyme activity was higher in isolated ECs from FASi-treated diabetic controls compared to all other groups **(Figure 3F-I).** Glucose and lipid uptake were decreased while lipid oxidation increased in ECs from FASi-treated diabetic controls compared to FASi-treated diabetic mice **(Figure 3J-K)**. FASi treatment in diabetic emut mice caused a significant increase in the lipid oxidation as measured by ^C14^CO_2_ release in isolated ECs compared to that in vehicle-treated diabetic emut mice **(Figure 3K)**. Taken together these data suggest that ANGPTL4 regulates de novo lipid biogenesis in diabetic endothelium. We further investigated the effects of a glycolysis inhibitor (2-DG) and a PKM2 inhibitor (PKM2i) in diabetic mice. 2-DG and PKM2 activator (TEPP-46) did not cause any alteration in blood glucose but significantly reduced the levels of ACR and relative collagen in diabetic mice **(Figure S6)**.

### Endothelial ANGPTL4-depleted mice suppress c-GAS-STING pathways in diabetic endothelium

De novo lipogenesis and increased lipid uptake leads to lipid accumulation in diabetic ECs^93,102,103^. This lipid accumulation along with the accumulation of glycolysis intermediates are detrimental to cell organelles such as mitochondria^82,83,85,103,104^. Damaged mitochondria release mitochondrial DNA in the cytosol that is sensed by the cytosolic cGAS-stimulator of interferon genes (STING) DNA sensing pathway (c-GAS-STING pathway), which subsequently upregulates the transcription of proinflammatory genes^59,94^. The mitochondrial marker protein TFAM was suppressed in CD31-positive cells isolated from kidneys of diabetic control mice compared to cells from nondiabetic control kidneys. TFAM levels in CD31-positive cells were substantially increased in cells from diabetic emut mice compared to cells from diabetic controls **(Figure 4A)**. In isolated ECs from diabetic control mice, expression of the mitochondrial genes TFAM and NDUFV2 was down-regulated, while expression of the c-GAS gene TMEM173, the STING gene Mb21d2 and inflammatory genes such as IL-1β, IL-6 and was upregulated **(Figure 4B)**. Isolated ECs from diabetic emut mice had elevated mitochondrial gene expression and suppressed expression of TMEM173, Mb21d2, IL-1β, IL-6 and NFkB1 **(Figure 4B)**. Isolated cytosolic fractions were then obtained from nondiabetic and diabetic mice. ECs from diabetic mice had increased expression of the mitochondrial genes mt-Co1 and mt-Cyb in the cytosolic fraction compared to non-diabetic ECs. ECs from emut mice had significantly suppressed expression of mt-Co1 and mt-Cyb in cytosolic fractions **(Figure 4C)**, suggesting mitochondrial DNA release and c-GAS-STING activation in diabetic ECs from control mice: this expression was decreased in ANGPTL4-depleted diabetic endothelium Immunofluorescent staining revealed that expression of phospho-IKBα in isolated ECs and CD31-positive cells from kidneys of diabetic controls was increased compared to the much lower expression noted in isolated ECs and CD31-positive cells from kidneys of diabetic emut mice **(Figure 4D; Figure S7A)**. Similarly, nuclear p-65 expression levels were higher in kidneys and isolated ECs from diabetic controls while relatively suppressed in tissues and ECs from diabetic emut mice **(Figure 4E; Figure S7A)**. Nondiabetic mice of each genotype showed no differences in expression of either p-IKBα or p-65 **(Figure S7A)**. Gene silencing of TMEM173 and Mb21d2 resulted in reduction of phospho-IKBα, and nuclear p-65 expression as well as the gene expression levels of *IL-1*β, *IL-6* and *NFKB1* in ECs **(Figure S7B-C)**. We observed renal IL-1β, IL-6, and MCP-1 levels were significantly higher in diabetic controls compared to nondiabetic mice, and these levels were significantly lower in diabetic emut mice compared to diabetic controls **(Figure S8)**.These data suggest that Angptl4 deficiency in endothelium suppresses diabetes-associated cytokines, mtDNA release and c-GAS-STING activation.

**Figure 4:**
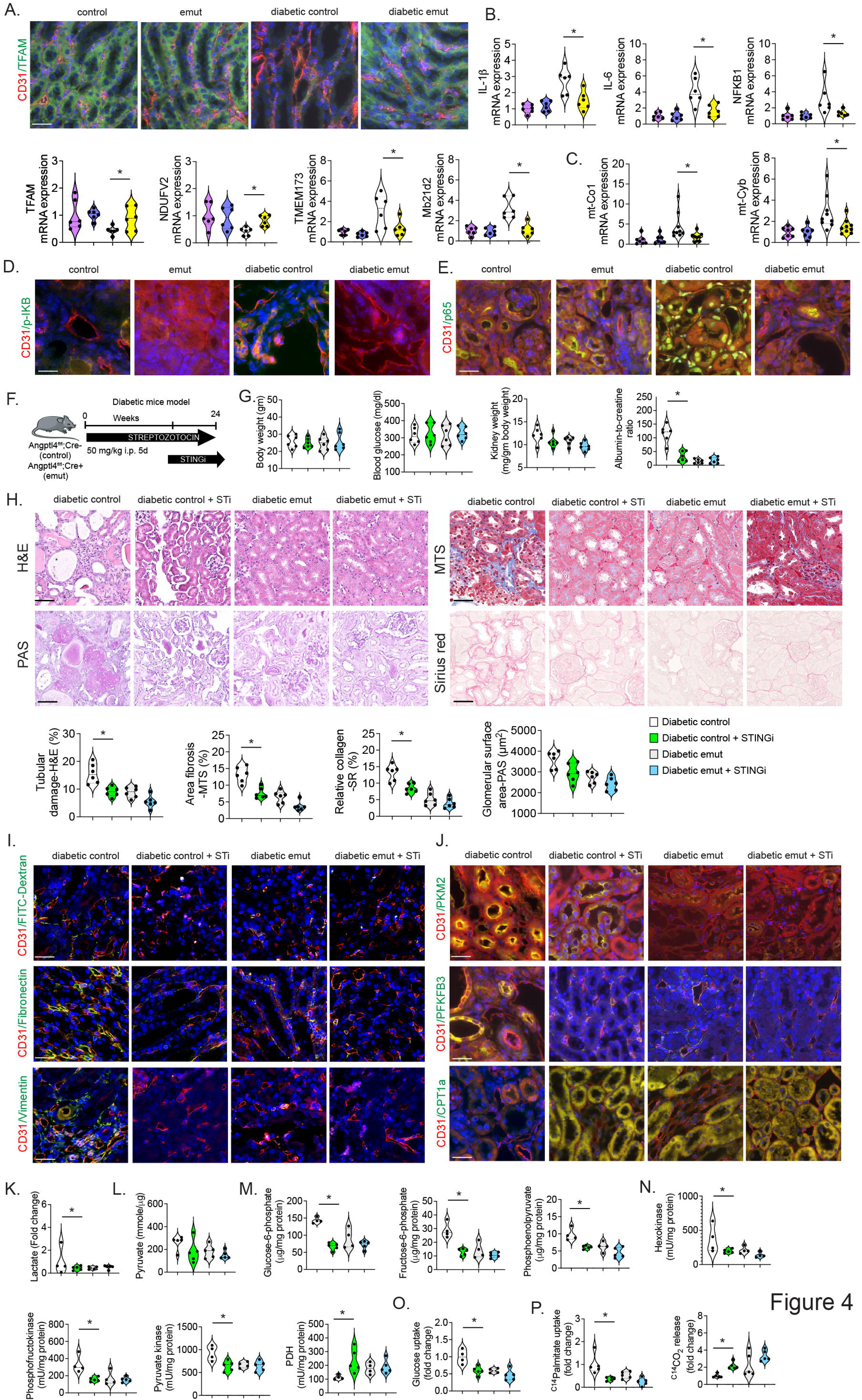
STING inhibition protects against DKD in diabetic control mice but not in diabetic emut mice. **A.** Immunofluorescence analysis of CD31/TFAM in the kidneys of nondiabetic and diabetic control and emut mice. FITC-TFAM; Rhodamine-CD31 and DAPI-blue. Representative images are shown. N=6/group. Original magnification 40x. Scale bar 50 μm. **B.** Relative mRNA expression levels of the indicated genes in isolated ECs from kidneys of the indicated mice. 18S was used as an internal control to normalize the data. **C.** Relative gene expression levels of mitochondrial genes *mt-Co1* and *mt-Cyb* in the cytosolic fractions of ECs. N = 8/group 18S was used as an internal control. **D.** Immunofluorescence analysis of p-IKB/CD31 in the kidneys of nondiabetic and diabetic control and emut mice. N = 6/group. FITC-p-IKB; rhodamine-CD31, and DAPI-blue. Representative images are shown. Original magnification 40X. Scale bar, 50 μm. N = 6 mice/group. **E.** Immunofluorescence analysis of p-65/CD31 in kidneys N = 6/group. FITC-p-65; rhodamine-CD31, and DAPI-blue. Representative images are shown. Original magnification 40X. Scale bar, 50 μm. N = 6 mice/group. **F.** A schematic showing the treatment protocol using the STING inhibitor (STINGi). **G.** Body weight, blood glucose, kidney weight/body weight and albumin-to-creatinine ratio. N=6/group. **H.** Kidney histology was analyzed in the kidneys of STINGi treated diabetic control and emut mice. Representative images are shown. Tubular damage was analyzed from H&E, relative area of fibrosis was analyzed from MTS, relative collagen was analyzed from SR and glomerular surface area was measured from PAS images. Original magnification 40x. N=6/group Scale bar 50 μm. **I.** Endothelial permeability was measured by FITC-dextran by immunofluorescence FITC-FITC-dextran, Rhodamine-CD31 and DAPI-blue. Original magnification 40x. N=6/group. Representative images are shown. Scale bar 50 μm. **J.** Immuno-co-labelling of CD31/Vimentin, CD31/Fibronectin, CD31/PFKFB3, CD31/PKM2, CD31/CPT1a in the kidneys of the STINGi treated diabetic control and emut mice. FITC-Vimentin, Fibronectin, PFKFB3, PKM2, CPT1a; Rhodamine-CD31; DAPI-blue. Representative images are shown. N=6/group. Original magnification 40x. Scale bar, 50 μm. **K.** Lactate measurement, **L.** pyruvate measurement, **M.** Glucose-6-PO_4_, fructose-6-PO_4_, phosphoenolpyruvate, **N.** hexokinase activity, phosphofructokinase activity, pyruvate kinase activity and pyruvate dehydrogenase enzyme activity measured in isolated ECs N=4-5/group **O.** Glucose uptake assay in isolated ECs using the calorimetric method. N=5/group **P.** Lipid uptake and lipid oxidation were measured as ^14C^palmitate uptake and ^14C^palmitate oxidation using established methods in the indicated groups. CPM was counted using a scintillation counter. Data are presented as mean±SEM. One-way-Anova Tukey’s test was used to calculated statistical significance among the groups.

To investigate the STING pathway and inflammation in diabetic endothelium, we used a small molecule inhibitor of STING (STINGi-C-176) and treated diabetic mice for 8-weeks followed by analysis of physiological parameters and kidney histology **(Figure 4F)**. There was no remarkable change in body weight, blood glucose or kidney weight between diabetic groups, however we found significant suppression in ACR in the STINGi-treated control mice compared to vehicle-treated control mice **(Figure 4G)**. We found a significant reduction in tubular damage, area of fibrosis and collagen deposition in STINGi-treated diabetic control mice. We did not observe any remarkable changes in tubular damage, area of fibrosis, collagen deposition or glomeruloscelrosis in treated or untreated emut mice **(Figure 4H)**. Next, the phenotype of the STINGi-treated mice was evaluated in the accelerated UUO model^97^. In this model, STINGi was provided to UUO operated-control and emut mice were then subjected to 4 daily doses of streptozotocin and euthanasia on day 11. STINGi-treated control mice had lessened fibrosis compared to UUO-operated control mice; no differences were observed in the emut mice **(Figure S9)**.

Endothelial cell leakage was remarkably suppressed in STINGi-treated diabetic mice compared to diabetic controls, while no substantial change was noted in emut mice **(Figure 4I)**. The kidneys of STINGi-treated diabetic mice had suppressed EndMT and fibrosis as evidenced by lower levels vimentin and fibronectin staining in CD31-positive cells which was associated with reduced levels of PFKFB3 and PKM2 and increased expression of CPT1a compared to CD31-positive cells from vehicle-treated diabetic mice **(Figure 4J)**. Lactate, pyruvate, glucose-6-PO_4_, fructose-6-PO_4_, phosphoenolpyruvate and glycolysis enzymes such as hexokinase, phosphofructokinase and pyruvate kinase were lower while PDH enzyme activity was higher in isolated ECs from STINGi-treated diabetic controls compared to those from vehicle-treated diabetic control mice **(Figure 4K-N).** Glucose uptake and lipid uptake were lower while lipid oxidation was higher in ECs from STINGi-treated diabetic controls compared to vehicle-treated diabetic mice **(Figure 4O-P)**. STINGi-treatment in diabetic emut mice caused a significant increase in lipid oxidation in isolated ECs compared to vehicle-treated diabetic emut mice **(Figure 4P)**. Taken together these data suggest that ANGPTL4 depletion in the endothelium suppresses lipotoxicty/glucotoxicy-related mitochondrial damage and c-GAS-STING activation-mediated inflammation in diabetes.

### Endothelial ANGPTL4 depletion augments VEGFR1 to VEGFR2 switching in diabetic endothelium leading to protection from DPP-4-mediated mesenchymal activation in tubules

VEGFR2 to VEGFR1 switching is the key phenotype associated with mesenchymal activation in diabetic ECs^100,105^. We found increased VEGFR1 and decreased VEGFR2 in CD31-positive cells from kidneys of diabetic control mice compared to cells from non-diabetic mice. Kidneys from diabetic emut had suppressed VEGFR1 and induced VEGFR2 in CD31-positive cells, suggesting VEGFR1-to-VEGFR2 switching is regulated by ANGPTL4 **(Figure 5A).** VEGFR1-to-VEGFR2 switching is associated with decreased expression of DPP-4 and β1-integrin in CD31-positive cells in kidneys from diabetic emut mice compared to those from diabetic control mice **(Figure 5B).** To investigate the role of DPP-4 and β1-integrin in the initiation of mesenchymal activation (partial EMT) in the tubules, the conditioned media from isolated ECs from diabetic mice of both genotypes was transferred to healthy isolated tubules **(Figure 5C).** Exposure to the conditioned media from diabetic controls led to a gain of α-SMA, DPP-4 and smad3 phosphorylation in tubules while the conditioned media from diabetic emut resulted in suppression of these 3 read-outs, suggesting partial mesenchymal activation in tubules is suppressed in the absence of ANGPLT4 **(Figure 5D).** This partial EMT process in diabetic controls is linked with higher in-situ proximity ligation of TGFβR1 and TGFβR2 dimerization and increased interaction between DPP-4 and β1-integrin; these processes are suppressed in diabetic emut mice **(Figure 5E).**

**Figure 5:**
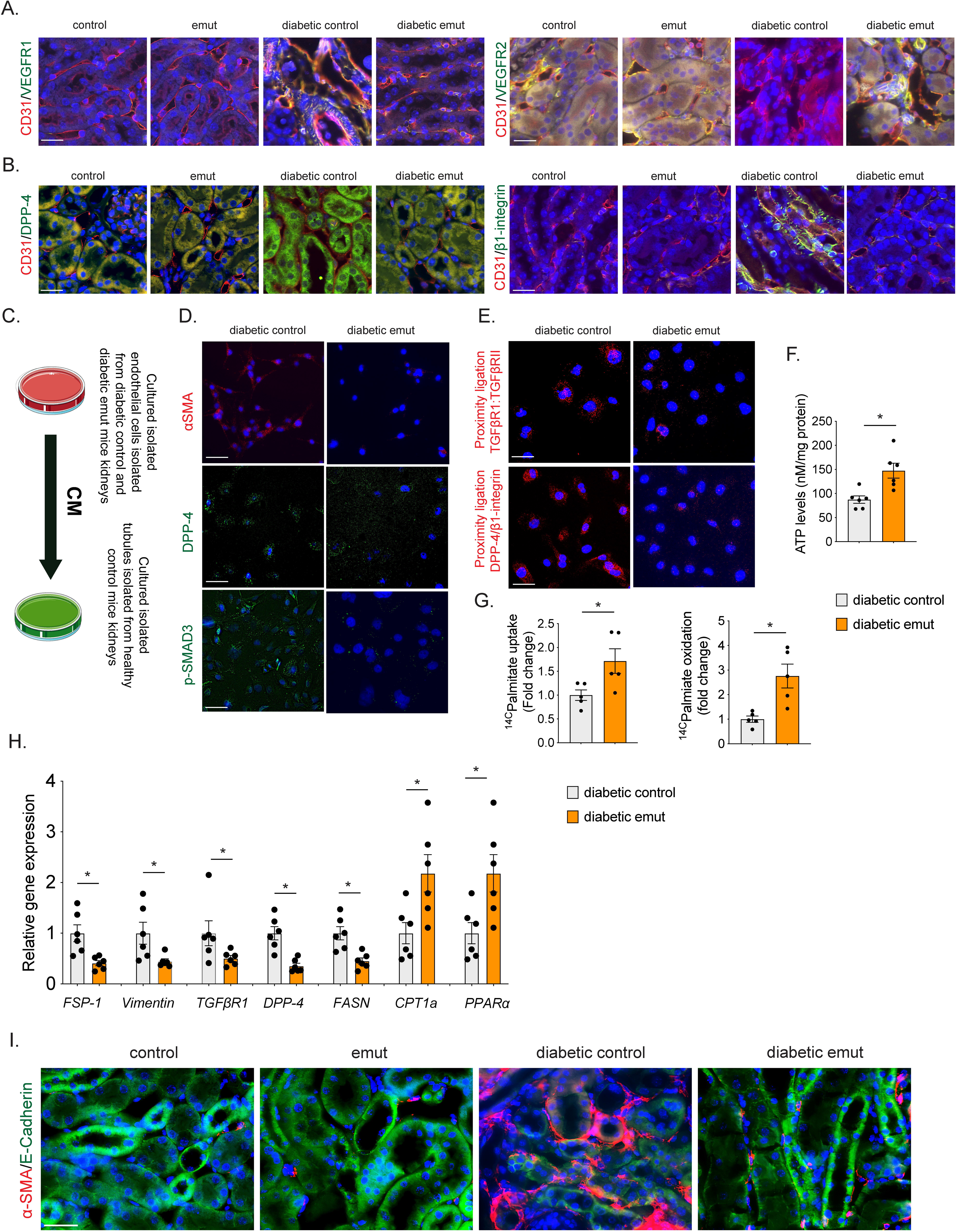
ANGPTL4 depletion mediates VEGFR1-to-VEGFR2 switching in diabetic ECs and, leads to suppression in DPP-4/β1-integrin related partial mesenchymal transition in tubules. A. Immunofluorescence analysis of CD31/VEGFR1 and CD31/VEGFR2 in kidneys of nondiabetic and diabetic control and emut mice. FITC-labeled-VEGFR1, VEGFR2; Rhodamine labeled-CD31 and DAPI-nucleus blue. Representative images are shown. N=6/group. Original magnification 40x. Scale bar 50 μm. **B.** Immunofluorescence analysis of CD31/DPP-4 and CD31/β1-integrin in kidneys of nondiabetic and diabetic control and emut mice. FITC-DPP-4, β1-integrin; Rhodamine-CD31 and DAPI-blue. Representative images are shown. N=6/group. Original magnification 40x. Scale bar 50 μm. **C.** Conditioned media (CM) experimental design. Three independent sets of experiments were analyzed**. D.** Immunofluorescence analysis of αSMA expression, DPP-4 and p-SMAD3 in conditioned media-treated TECs. For each slide, images of six different fields of view at 400x magnification were evaluated. FITC-DPP-4 and p-smad3l Rhodamine-αSMA; DAPI-blue. N = 3 independent samples/group. Scale bar, 30 μm. **E.** TGFβRI:TGFβRII and DPP-4:β1-integrin proximity was analyzed by Duo-link proximity ligation assay in isolated cells of CM media-treated isolated tubules (from diabetic control and diabetic emut). Representative images are shown. Three independent sets of experiments were analyzed. For each slide, images of six different fields of view at 400x magnification were evaluated. Scale bar 30 μm. **F.** Cellular ATP measurement. Six independent samples were analyzed. **G.** [^14^C]palmitate uptake and [14C]palmitate oxidation as measured by [^14^CO_2_]release. CPM was counted and normalized to the protein in the well. N = 5 independent samples/group. **H.** Relative mRNA levels of *FSP-1, Vimentin, TGFβR1, DPP-4, FASN, CPT1a, PPARα* determined by qRT-PCR in conditioned media-treated isolated TECs. N = 6 independent replicates per group. Data are presented as mean ± SEM. Student’s t test was used for the analysis of statistical significance.

In cultured HK-2 cells, TGFβ1 treatment increased the interaction between ANGPTL4 and β1-integrin **(Figure S10A).** Simultaneous treatment of HK-2 cells with TGFβ1 and either DPP-4 siRNA or β1-integrin siRNA substantially suppressed the proximity interactions between ANGPTL4 and β1-integrin **(Figure S10B)**. ATP and FAO levels were significantly higher in diabetic emut mice compared to diabetic controls **(Figure 5F-G).** Relative gene expression levels of FSP-1, Vimentin, TGFβR1, DPP-4, and FASN were significantly down-regulated while CPT1a and PPARα were significantly up-regulated in diabetic emut mice compared to diabetic controls, supporting the hypothesis that endothelial ANGPTL4 depletion results in VEGFR1-to-VEGFR2 switching in diabetic endothelium, and leads to suppression of DPP-4/β1-mediated mesenchymal activation in tubules, suppression of EndMT and preservation of metabolic homeostasis **(Figure 5H).**

### Endothelial ANGPTL4 depletion results in SIRT1 induction and SIRT1 overexpression in endothelium is protective in DKD

To investigate further the consequences of endothelial ANGPTL4-loss in diabetes, we used previously published NEXT-generation sequencing data and compared the gene signatures in ANGPTL4-siRNA and control-siRNA transfected HUVECs. We observed the Sirtuin1 (SIRT1) level was higher in the ANGPTL4-siRNA transfected group. Elevated SIRT1 has been associated with lipid and glucose metabolism and mesenchymal transition in other cell types^29^ **(Figure S11)**. We confirmed SIRT1 expression levels by immunofluorescence in our mouse models and observed that, in general, diabetic endothelium had suppressed expression of SIRT1 which was most pronounced in the diabetic control kidneys **(Figure S12A)**. Diabetic emut mice showed some increase in staining intensity suggesting that ANGPTL4 loss directly or indirectly associates with SIRT1 induction in CD31-positive cells **(Figure S12A)**. We then analyzed the expression of several known anti-mesenchymal and anti-fibrotic molecules including endothelial SIRT3, endothelial glucocorticoid receptor (GR), and endothelial FGFR1, all of which have been shown to be effective in mouse models of diabetes^35,81,98,99,106^. We found all three molecules demonstrated increased expression in kidneys from diabetic emut mice compared to kidneys from diabetic control mice **(Figure S12B)**.

To further investigate the role of SIRT1 in diabetic endothelium we generated an endothelial SIRT1 overexpression mouse (eEx-SIRT1) which was on the C57B/L6 background **(Figure 6A; Figure S12C-D)**. Control and eEx-SIRT1 mice were injected with streptozotocin and provided with free access to food and water for 16 weeks. Then they were sacrificed and physiological parameters and kidney histology were analyzed **(Figure 6A)**. We did not observe any changes in body weight, blood glucose or kidney weight, however diabetic eEX-SIRT1 mice had suppressed ACR compared to diabetic controls **(Figure 6B)**. Kidney histology demonstrated suppressed tubular damage, area of fibrosis, collagen deposition and glomerulosclerosis in kidneys of diabetic eEx-SIRT1 mice **(Figure 6C)**. Vimentin and fibronectin levels were reduced in CD31-positive cells from kidneys of diabetic eEx-SIRT1 mice compared to levels in diabetic control mice. PFKFB3 and PKM2 levels were suppressed while CPT1a levels were induced in CD31-positive cells from kidneys of diabetic eEx-SIRT1 mice compared to kidneys from diabetic controls **(Figure 6D)**. Lactate levels and glycolysis intermediates including pyruvate, glucose-6-PO_4_, fructose-6-PO_4_, phosphoenolpyruvate and glycolysis enzymes such as hexokinase, phosphofructokinase and pyruvate kinase were higher while PDH enzyme activity was lower in isolated ECs from diabetic controls. The opposite pattern was observed in isolated ECs from diabetic eEx-SIRT1 mice **(Figure 6E-H).** Glucose uptake was higher in ECs from diabetic controls, while suppressed glucose uptake, similar to the non-diabetic mice, was observed in isolated ECs from diabetic eEx-SIRT1 mice **(Figure 6I).** Taken together these data demonstrate that ANGPTL4 depletion restores SIRT1 levels and SIRT1 overexpression in endothelium improves associated defects in lipid and glucose metabolism in diabetes.

**Figure 6:**
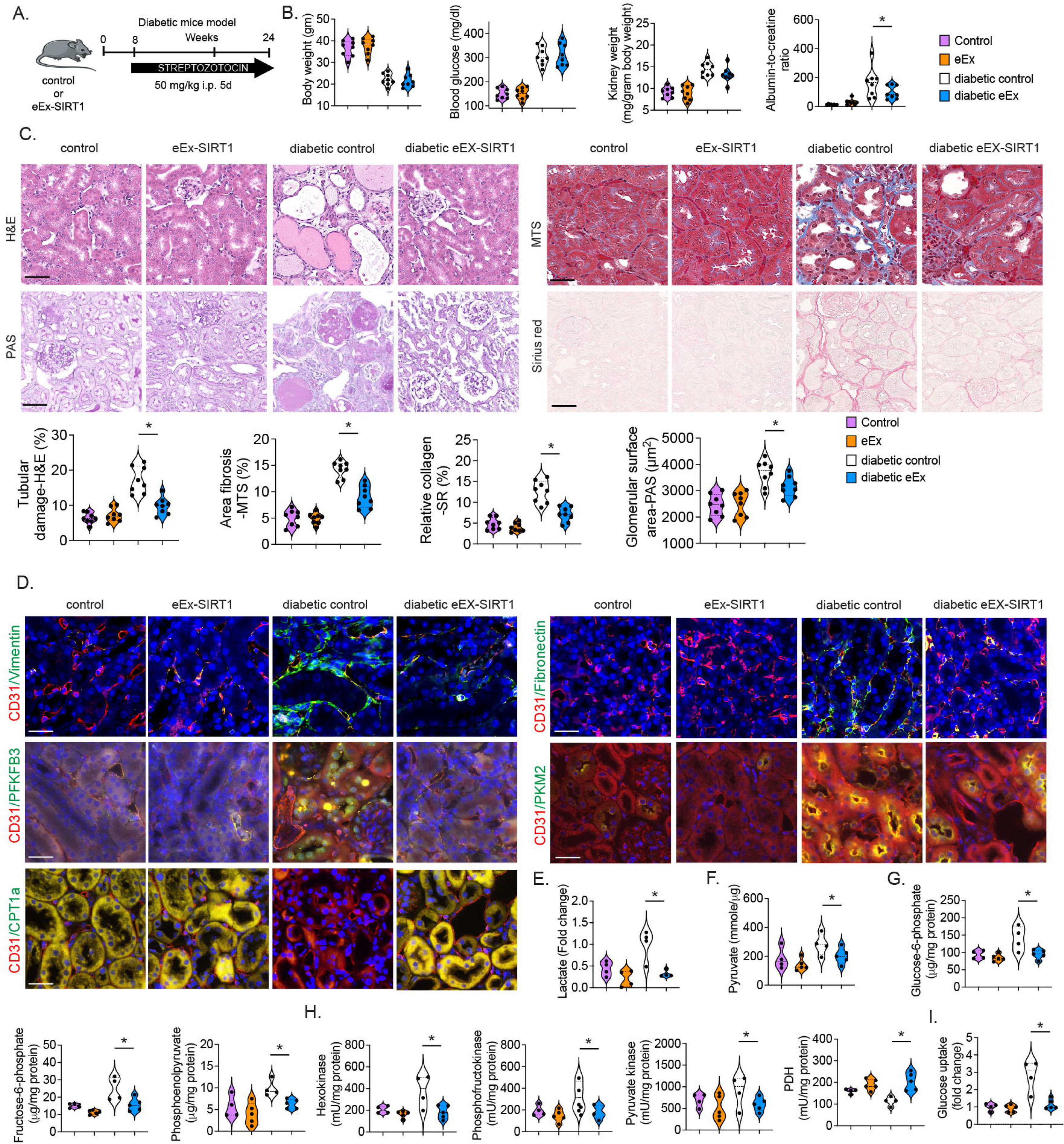
Endothelial SIRT1 overexpression protects against DKD. **A.** Schematic showing diabetes induction in control and endothelial SIRT1 overexpression mice. **B.** Body weight, blood glucose, kidney weight/body weight and urine albumin-to-creatine ratio. N=8/group. **C.** Histology in kidneys from the indicated mice. Representative images are shown. Tubular damage was analyzed from H&E, relative area fibrosis was analyzed from MTS, relative collagen was analyzed from SR and glomerular surface area was measured from PAS images. Original magnification 40x. N=8/group. Scale bar 50 μm. **D.** Immuno-co-labelling of CD31/Vimentin, CD31/Fibronectin, CD31/PFKFB3, CD31/PKM2, CD31/CPT1a were analyzed in kidneys from the indicated mice. FITC-Vimentin, Fibronectin, PFKFB3, PKM2, CPT1a; Rhodamine-CD31; DAPI-blue. Representative images are shown. N=6/group. Original magnification 40x. Scale bar 50 μm. **E.** Lactate measurement, **F.** pyruvate measurement, **G.** Glucose-6-PO_4_, fructose-6-PO_4_, phosphoenolpyruvate, **H.** hexokinase activity, phosphofructokinase activity, pyruvate kinase activity and pyruvate dehydrogenase enzyme activity measured in isolated ECs from the kidneys. N=4-5/group. **I.** Glucose uptake assay in isolated ECs using the calorimetric method. N=5/group. Data are presented as mean ± SEM. One-way-Anova Tukey’s test was used to calculated statistical significance among the groups.

## Discussion

Both paracrine and autocrine roles of endothelial-cell ANGPTL4 are largely unknown, especially in the renal vasculature in the setting of diabetes mellitus. Here we report that endogenous ANGPTL4 promotes a metabolic phenotype which favors mesenchymal activation in ECs and in renal tubules, which leads to fibrogenesis in diabetic kidneys. We demonstrate that diabetic kidney fibrosis is associated with higher EC-ANGPTL4 gene expression which regulates local lipase, and induces fatty acid uptake, glucose uptake and abnormal glycolysis The cumulative result of these effects is lipotoxicity and glucotoxicity in ECs, which leads to aberrant angiogenesis and inflammation in diabetic kidneys. Depletion of EC-ANGPTL4 mitigates lipotoxicy, glucotoxicity and inflammation and suppresses related mesenchymal activation and fibrogenesis in DKD.

ANGPTL4 has shown diverse functions and its expression is largely influenced by tissue microenvironments and pathological conditions. Human studies have shown a positive correlation between Angptl4 levels and expression of characteristics of diabetic nephropathy^107^. In cultured glomerular mesangial cells, Angptl4 deficiency inhibits both the inflammatory response and extracellular matrix accumulation^108^. These observations implicate Angptl4 as a potential therapeutic target for the treatment of DKD. However, a clear role for Angplt4 in regulating whole body lipid and glucose metabolism has not been fully elucidated due to lack of availability of relevant tissue-specific knock out mouse models. Severe systemic metabolic complications such as gut inflammation and ascites have limited investigators’ ability to analyze the beneficial functions of Angptl4 deficiency in diabetes^109^. Most of the angiogenic studies related to the ANGPTL4 have been done using whole-body knockout mice or in vitro cell culture systems and have demonstrated conflicting results^67,69,72,74,110^. Therefore we utilized endothelial-specific ANGPTL4 to clearly decipher its role in renal angiogenesis and its relationship to fibrogenic processes.

*Angptl4* is a known inhibitor of LPL, an enzyme that catalyzes the hydrolysis of triglycerides into fatty acids, which are utilized by peripheral tissues and kidney tubules^111^. ANGPTL4 also inhibits endothelial lipase in ECs^29,57,112^. In our previous study we demonstrated that *Angptl4* expression is upregulated, and LPL expression and its associated activities are downregulated, in fibrotic kidneys during diabetes. However, while chemical inhibition of LPL in diabetic mice caused an elevation in plasma triglycerides, it did not alter renal fibrosis, suggesting that LPL is not a key molecule in the development of diabetic kidney fibrosis. Here, we used a small molecule inhibitor specific for endothelial lipase and treated diabetic mice. However, we did not observe any remarkable effects of this inhibitor on fibrogenesis, suggesting that endothelial lipase is not a key molecule associated with diabetic kidney fibrosis. Analysis of diabetic kidneys from endothelial Angptl4 mutant mice showed suppression of renal fibrosis, proteinuria, pro-inflammatory cytokines, and mesenchymal activation in tubules and ECs when compared to diabetic controls, suggesting that endothelial *Angptl4* is pro-fibrotic and pro-inflammatory in diabetes. Our data clearly demonstrate that endothelial Angptl4 is one of the catalysts of renal fibrosis in diabetes and leads to disruption of cytokine and chemokine metabolic reprogramming by up regulating DPP-4/β1-integrin-associated TGFβ signaling. These processes alter metabolic homeostasis by predisposing towards defective fatty acid metabolism and aberrant glucose metabolism-associated mesenchymal activation in ECs and in neighboring tubules. Our results further emphasize that endothelial ANGPTL4 is a required protein to support pathological angiogenesis in diabetic kidneys. Although physiological and pathological angiogenesis shares many common characteristics, some distinctions between two processes imply subtle differences in underlying mechanisms. The interplay of endothelial ANGPTL4 with different known regulators of angiogenesis in physiological vs. pathological settings is an understudied area that warrants further investigation.

Clement and colleagues demonstrated the critical role of podocyte-specific Angptl4 in causing proteinuria in a mouse model of nephrotic syndrome^75^. Angplt4 was reported to express in two forms, the sialylated (normal) form and the hyposialylated (abnormal) form. The hyposialylated form is causative for proteinuria. Conversion of the hyposialylated to the sialylated form by treatment with N-acetyl-D-mannosamine (NAM) was noted to significantly suppress the levels of proteinuria in a mouse model of nephrotic syndrome^75^. Clement et al., also demonstrated that circulating sialylated Angplt4 suppresses proteinuria while increasing hypertriglyceridemia in mice^76^. In line with previous findings, our previous data demonstrate that N-acetyl-D-mannosamine treatment in diabetic mice suppressed fibrosis and proteinuria and restored kidney structure in diabetic mice^59^. Further, our data show that blocking Angptl4 in ECs effectively suppresses proteinuria, glomerular fibrosis and interstitial fibrosis in diabetic mice. One limitation of our study is that we could not demonstrate whether endothelial Angptl4 in diabetic kidneys exists in the hyposialylated or sialylated form due to the lack of a suitable mouse Angplt4 antibody, thus precluding us from evaluating endothelial Angptl4 at the protein level. However, based on previous and current data, we suspect that diabetes may cause endothelial secretion of the hyposialylated form of Angptl4 in the kidney.

To investigate how endothelial Angptl4 accelerates mesenchymal activation, we analyzed lipid levels, glycolysis metabolites and FAO in ECs. It is clear from our results that in diabetic endothelium, Angptl4 has a prominent effect on de novo lipogenesis (DNL). The elevated triglycerides (lipid load) and glycolysis metabolites due to defects in FAO in mitochondria and aberrant glycolysis were suppressed by depletion of endothelial Angptl4 which reduced the lipid and glycolysis metabolite loads, restored mitochondrial function and restored lipid oxidation in diabetes. It has been reported that increased FAO as a result of overexpression of Cpt1a improves mitochondrial homeostasis and fibrosis in mice^103^ and lipid accumulation and DNL have been shown to be higher in the kidneys of patients with fibrosis^93^. In addition, such abnormal DNL is regulated by Acyl-CoA synthetase short-chain family 2, leading to NADPH depletion and increased ROS levels, ultimately promoting NLRP3-dependent pyroptosis in kidney tubules^93^. These processes are key to the induction of myofibroblast activation in diabetic endothelium and fibrosis in diabetic kidneys^82,93^. Our data suggest that upregulation of *Angptl4* expression in diabetic endothelium is associated with increased fibrosis, lipid accumulation and glycolysis and suppression of FAO, leading to a heightened angiogenic response, Endothelial *Angptl4* deficiency results in increased lipid oxidation, decreased abnormal glucose metabolism and diminished lipotoxicity, suggesting that metabolic reprogramming by *Angplt4* deficiency is a crucial step in promoting anti-mesenchymal and anti-fibrotic mechanisms in renal ECs and tubules during diabetes.

Metabolic shifts are critical for mesenchymal transformation in ECs and related extracellular matrix deposition in diverse type of cells in the kidney^25,83,85,86^. Quiescent ECs mostly rely on FAO for ATP generation while stimulated angiogenic ECs use glycolysis to support their growth and proliferation to form vessel-like structures^81^. Therefore, growing ECs are dependent on the glycolysis-tricarboxylic acid (TCA) cycle for energy and proflieration^81^. Our results demonstrate that kidney ECs undergo shifts in the metabolic fuel from FAO to abnormal glycolysis, and these metabolic shifts are more prominent in diabetic controls. This abnormal glycolysis is different from the normal glycolysis process that terminates in the TCA and electron transport chain (ETC) cycles^89^. This abnormal glycolysis-lactate cycle terminates in lactate production which has been found to accelerate fibrotic processes in many cell types^113–116^. It is evident from our results that higher levels of lactate, pyruvate and glycolytic metabolites exist in diabetic controls, suggesting that metabolic shifts from normal to abnormal glycolysis contributes to extra-cellular matrix (ECM) production in diabetic ECs. This defective FAO and aberrant glycolysis is reversed in EC-specific ANGPTL4 mutant mice, suggesting that renal endothelial ANGPTL4 contributes to DNL, defective lipid oxidation and aberrant glycolysis-lactate cycles. It is assumed that lactate derived from diabetic ECs induces fibrotic signaling and mesenchymal transdifferentiation processes in other kidney cell types, most notably tubules, and this partial EMT is the one of the established processes in the development of kidney fibrosis in diabetes^8,9,20^.

Lipotoxicity and glucotoxicity and related mitochondrial defects in diabetic endothelium are crucial in kidney disease development^35,81,82,103^. These abnormal metabolic processes contribute to cellular energy deficits, compromised EC function and lack of proper mitochondrial integrity which are, in turn, linked with release of mtDNA into the cytoplasm of diabetic ECs, where it is recognized as “foreign” DNA. This recognition inappropriately activates the immune system, triggering pathological inflammation by activating the cytosolic cGAS-STING DNA sensing pathway resulting in aberrant cytokine expression and immune cell recruitment^94,117^. Blocking this foreign DNA sensor pathway in the endothelium mitigates development of DKD and could offer a new therapeutic strategy for patients with chronic kidney disease^94,118^. To determine how endothelial Angptl4 deficiency suppresses inflammation, we analyzed mtDNA release which was increased in ECs from diabetic control mice and significantly reduced in EC-specific Angptl4 mutant mice. Decreased mtDNA release and suppressed c-GAS-STING activation further decreased proinflammatory cytokine expression in diabetes suggesting that EC Angptl4 deficiency restores mitochondrial structure and function.

To understand how endothelial Angptl4 regulates lipid oxidation and defective glycolysis, we analyzed DPP-4-Integrin β1 signaling which is associated with TGFβR1 and TGFβR2 dimerization, the first crucial step in activating the TGFβRs-smad2/3 signalling cascade. Activated TGFβ signalling is known to suppress lipid oxidation and exacerbates abnormal glycolysis thereby activating ‘mesenchymal metabolic shifts’ in diabetic ECs and surrounding diabetic tubules^82,89^. DPP-4 is a well-known pro-EMT and pro-EndMT molecule in diabetic ECs^100,119,120^. It is evident from our results that TGFβ1-associated mesenchymal transformation is associated with closer proximity of β1-integrin in the presence of Angptl4 and with DPP-4 in diabetic ECs suggesting that Angptl4 interacts with Integrin β1 and influences activation of the DPP-4/β1-integrin mediated mesenchymal signal transduction cascade in diabetic ECs and tubules. Smad3 phosphorylation alters the cytoplasmic pathway and also enters the nucleus where it represses the transcription of FAO genes and induces the transcription of genes related to fibrogenesis in ECs and tubules^82^. These data indicate that endothelial Angptl4 interacts with β1-integrin and influences DPP-4/β1-integrin interactions and the associated TGFβ-smad3 signaling cascade, The cumulative effects of activation of these aberrant pathways lead to the activation of mesenchymal signal transduction in diabetic ECs and tubules.

Past studies have shown Angptl4 to be a potential metabolic regulator which is linked with several diseases including diabetes, atherosclerosis and cancer^121–123^. In a previous study, researchers used a neutralizing antibody to Angptl4 in both humanized mice and non-human primates which resulted in severe side effects and systemic metabolic abnormalities^80^. Our data using EC-specific Angptl4 mutant mice suggest that endothelial Angptl4 has promising drug targetability against DKD. Moreover, ANGPTL4 deficiency in ECs promotes known known anti-mesenchymal molecules such as SIRT3, FGFR1 and GR^35,81,90,98,99^. In addition, we demonstrated that endothelial SIRT1 levels were induced in the Angplt4-depleted condition, suggesting that the ANGPTL4 deficiency-associated elevation in SIRT1 partially contributes to the observed anti-mesenchymal activity in ECs and neighboring cell types.

When conditioned media from control and ANGPTL4-depleted ECs was transferred to cultured tubular epithelial cells (TECs) from control kidneys, we observed induction of mesenchymal markers and activation of TGFβ and the DPP-4-β1-integrin signalling cascade in the controls; these processes were reversed in the ANGPTL4-depleted ECs. These findings suggest that ANGPTL4 associated EndMT leads to mesenchymal activation in kidney tubules in diabetes. ANGPTL4-depleted ECs have lower levels of TGFβ-smad3 and DPP-4-β1-integrin signalling, which are associated with lower levels of plasma pro-inflammatory cytokines and induced levels of lipid oxidation. The effects of these metabolic changes result in the activation of mesenchymal transformation in diabetic ECs, which appears to exert paracrine effects on neighboring TECs.

In conclusion, endothelial Angpl4 is key pro-EndMT molecule and its depletion is associated with protection against DKD by metabolic reprogramming which is driven by mitigating DPP-4-β1-Integrin signalling and related TGFβ-smad3 signalling (Figure 7). These data add substantial information to the understanding of Angplt4 in EC biology and offer an exciting possibility as a new therapeutic option for the treatment of DKD.

**Figure 7:**
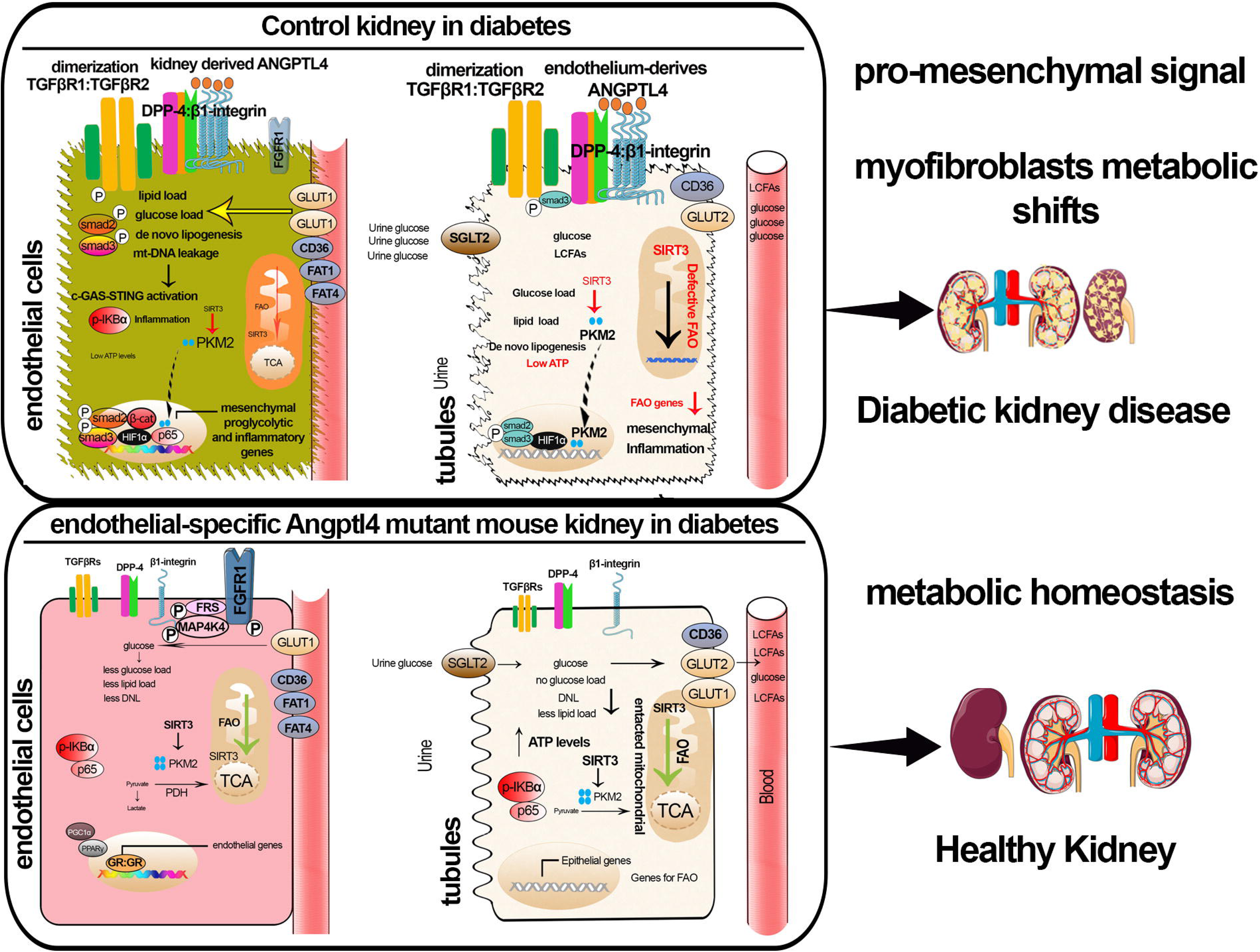
Schematic diagram summarizing the working hypothesis by which ANGPLT4 regulates EC metabolism in diabetes.

## Acknowledgements

Conceptualization: SS, AD, CFH, JG, DK, KK, Methodology: BA, AD, CFH, JG, DK, KK, GS, SS Investigation: SS, BA, LG, BLM, MM, GS, OS, RS, HZ

Validation: BA, KK Formal Analysis: BR, SS Visualization: AD, JG, SS

Supervision: AD, JG, DK, KI, KK, GS, SS Software: BR, SS

Data curation: SS Writing—original draft: SS

Writing—review & editing: AD, JG, SS

Resources: BA, TB, AD, CFH, JG, KK, BR, GS, SS

Funding Acquisition: SS, AD, JG, KI, KK, GS Project Administration: AD, JG, SS

---

**Figure S1: Validation of endothelial specific ANGPTL4 mutant mice**

**A.** β-gal staining in control and emut mice via immunofluorescence. N=5/group. FITC-β-gal, Rhodamine-CD31, DAPI-blue. Representative images are shown. Original magnification 40x. Scale bar 50 μm. **B**. Relative mRNA expression level of *Angptl4* analyzed by qPCR in isolated ECs from control and emut mice. 18S was used as an internal control. N=6/group. Data are shown as mean ± SEM. *p<0.05. Student-t test was used to calculated statistical significance among the groups

**Figure S2: Loss of endothelial ANGPTL4 protects against renal fibrosis in a mouse model of urinary obstruction (UUO).**

A. Masson trichrome staining (MTS) in contralateral and UUO-operated kidneys in control and emut mice were analyzed. Representative images are shown. Area of fibrosis (%) was measured using ImageJ. n=6/group. Data are shown as mean ± SEM. Scale bar 100 μm. *p<0.05. One-way-Anova Tukey’s test was used to calculated statistical significance among the groups

**Figure S3: Endothelial lipase inhibition did not cause any significant changes in diabetic mice**

**A.** Schematic showing the protocol for the treatment of XEN-445 in diabetic mice. **B.** Body weight, blood glucose, kidney weight/body weight (N=8/group) and urine albumin-to-creatine ratio (N=6/group). Blood glucose was analyzed using glucose strips. **C.** Histology was analyzed in kidneys from XEN-445-treated diabetic control and emut mice. Representative images are shown. Tubular damage was analyzed from H&E and relative collagen was analyzed from Siruis Red. Original magnification 40x. N=8/group. Scale bar 50 μm. Data are presented as mean±SEM. One-way-Anova Tukey’s test was used to calculated statistical significance among the groups.

**Figure S4. De novo lipogenesis in the kidneys of control and emut mice during diabetes**

**A.** Kidney triglycerides. N=5/group. **B.** Kidney total cholesterol levels. N=5/group.

**B.** qPCR gene expression analysis of *FASN*, *ACC* and *Perilipin, SREBF1, SCAP, CD36, FAT1, FAT4* in isolated ECs from the indicated mice. 18S was used as an internal control. D. CD31/FASN co-staining in kidneys. N=5/group. Representative images are shown. FITC-FASN, Rhodamine-CD31, DAPI-blue. Original magnification 40x. Scale bar 50 μm.

**Figure S5: Inhibition of fatty acid synthetase protects against renal fibrosis in a mouse model of urinary obstruction (UUO)**

A. Masson trichrome staining (MTS) in contralateral and UUO-operated kidneys from FASi-treated control and emut mice. Representative images are shown. Area of fibrosis (%) was measured using ImageJ. N=6/group. Data are shown as mean ± SEM. Scale bar 100 μm. *p<0.05. One-way-Anova Tukey’s test was used to calculated statistical significance among the groups

**Figure S6: Inhibition of glycolysis and activation of PKM2 protects against renal fibrosis in a mouse model of diabetes**

**A.** Schematic showing the protocol for the treatment of 2-DG and TEPP-46 (PKM2 activator) in diabetic mice. **B.** Blood glucose and urine albumin-to-creatine ratio. N=7/group. Blood glucose was analyzed using glucose strips.

**B.** Histology was analyzed in the kidneys from 2-DG and TEPP-46-treated diabetic controls. Representative images are shown. Relative collagen was analyzed from Sirius Red. Original magnification 40x. N=7/group. Scale bar 100 μm. Data in the graph are presented as mean±SEM. One-way-Anova and Tukey’s test was used to calculated statistical significance among the groups.

**Figure S7: c-GAS-STING knockdown suppresses inflammation in isolated ECs**

**A.** p-IKBα and p-65 staining in isolated ECs from the indicated conditions. Rhodamine-p-IKBα, FITC-p-65 and DAPI-blue. Original magnification 40x. Scale bar 50 μm. Experiments in triplicate were analyzed. Representative images are shown. **B.** p-IKBα and p-65 staining in TMEM173 siRNA- and Mb21d2 siRNA-transfected ECs. Experiments were performed in triplicate. Rhodamine-p-IKBα, FITC-p-65 and DAPI-blue. Original magnification 40x. Scale bar 50 μm. **C.** Relative gene expression analysis of *IL-1β, IL-6* and *NFkB* in isolated cells transfected with control siRNA, TMEM173 siRNA and Mb21d2 siRNA. 18S was used as an internal control. N=6/group. Data are shown as mean ± SEM. *p<0.05. One-way-Anova Tukey’s test was used to calculate statistical significance among the groups.

**Figure S8: Renal cytokine levels in mice**

A. Renal cytokines levels were analyzed by Luminex cytokine array. N=5/group. Data are shown as mean ± SEM. *p<0.05. One-way-Anova Tukey’s test was used to calculated statistical significance among the groups.

**Figure S9: STING inhibition protects against renal fibrosis in a mouse model of urinary obstruction (UUO).**

A. Masson trichrome staining (MTS) in contralateral and UUO-operated kidneys in STINGi-treated control and emut mice. Representative images are shown. Area of fibrosis (%) was measured using ImageJ. n=6/group. Data are shown as mean ± SEM. Scale bar 100 μm. *p<0.05. One-way-Anova Tukey’s test was used to calculated statistical significance among the groups

**Figure S10: ANGPTL4-**β**1-integrin proximity ligation assay**

**A.** ANGPTL4-β1-integrin proximity ligation assay in control and TGFβ1 (10ng/ml) stimulated HK-2 cells. Three independent experiments were analyzed. **B.** ANGPTL4-β1-integrin proximity ligation assay in TGFβ1 (10ng/ml)-treated cells transfected with either control SiRNA, DPP-4 siRNA or β1-integrin siRNAs. Three independent experiments were analyzed.

**Figure S11: Next-generation sequencing data**

**A.** Next-generation sequencing data in control siRNA- and ANGPTL4 siRNA-transfected HUVECs.

**Figure S12: SIRT3, FGFR1, GR and SIRT1 staining in isolated ECs**

**A.** Immunofluorescence analysis of CD31/SIRT3, CD31/FGFR1 and CD31/GR in kidneys from nondiabetic and diabetic, control and emut mice. N=6/group. FITC-SIRT3, FGFR1 and GR; rhodamine-CD31, and DAPI-blue. Representative images are shown. Original magnification 40X. Scale bar, 50 μm. **B.** Immunofluorescence analysis of CD31/SIRT1 in kidneys from the indicated mice. N=6/group. FITC-SIRT1; rhodamine-CD31, and DAPI-blue. Representative images are shown. Original magnification 40X. Scale bar 50 μm. **C.** qPCR gene expression analysis of SIRT1 mRNA in isolated ECs from control and SIRT1 overexpression mice (eEX-SIRT1). 18S was used as an internal control. N=8/group. **D**. Immunofluorescence analysis of SIRT1/DAPI in cultured isolated ECs from control and eEx-SIRT1 mice kidneys. FITC-SIRT1; DAPI-blue. Representative images are shown. Original magnification 40X. Scale bar 50 μm. Data are shown as mean ± SEM. *p<0.05. Student’s t test was used to calculated statistical significance among the groups

