## Supplementary material for "Metabolic reprogramming by endothelial ANGPTL4 depletion protects against diabetic kidney disease": Figures S1-S12

A.

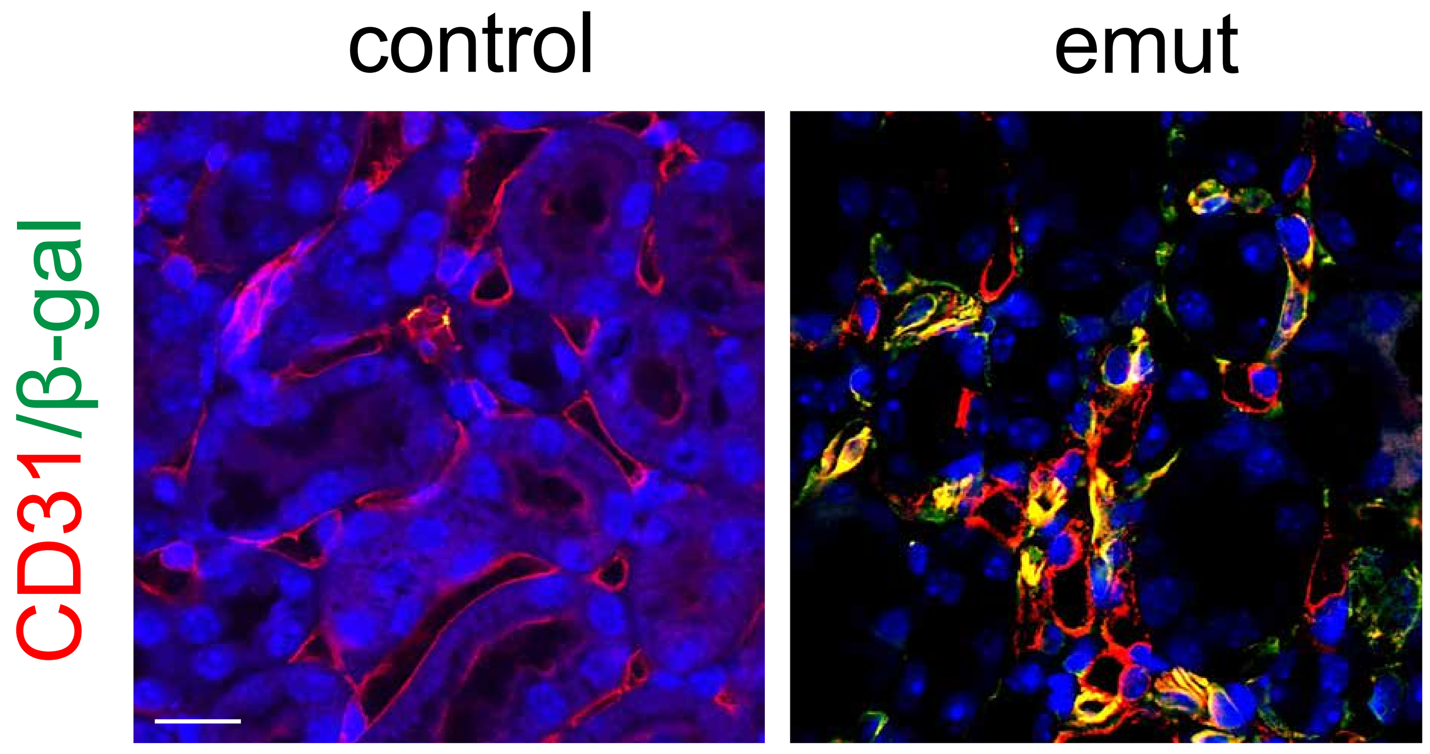
control
emut
CD31/β-gal

B.

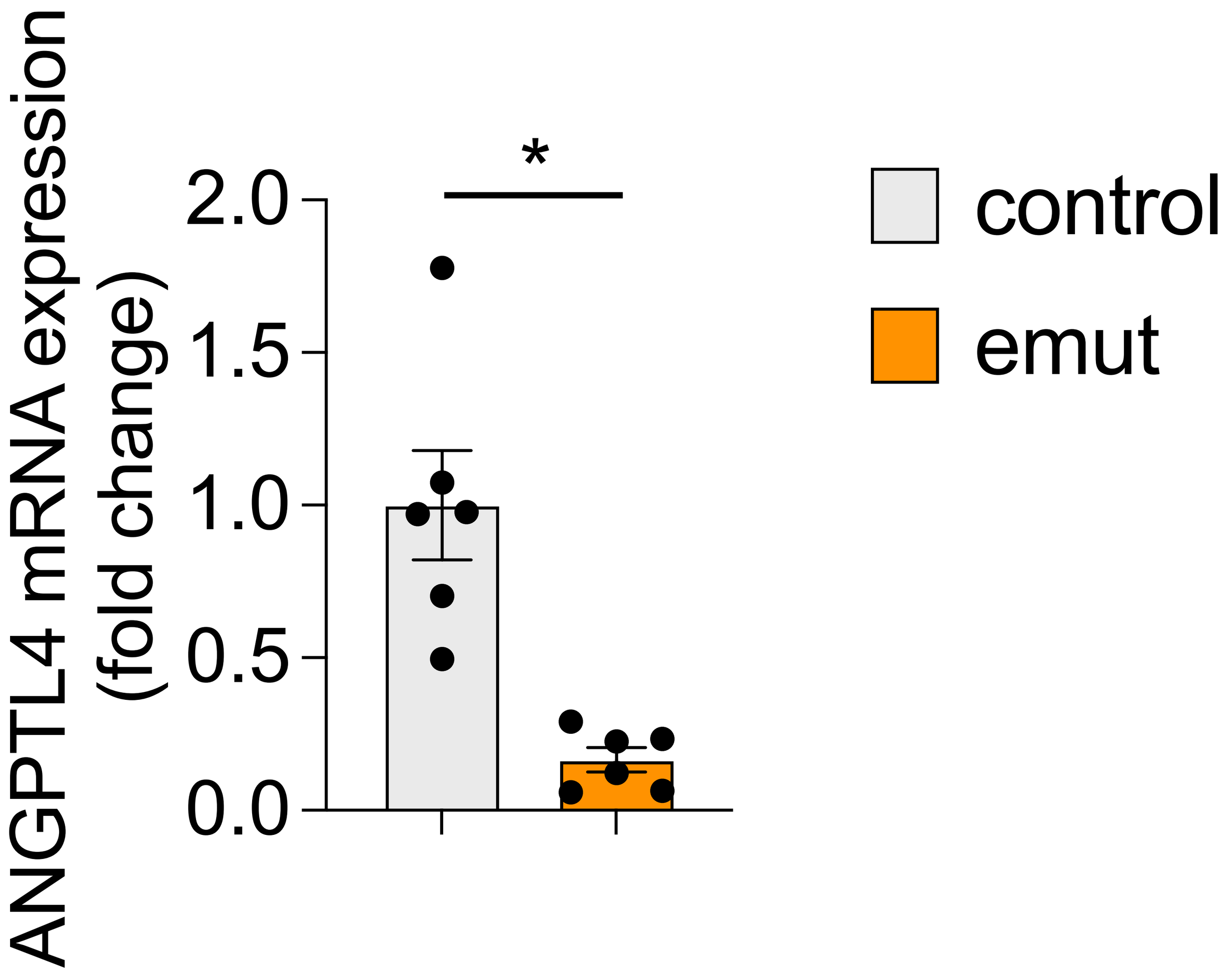
ANGPTL4 mRNA expression
(fold change)
2.0
1.5
1.0
0.5
0.0
*
control
emut

Figure S1

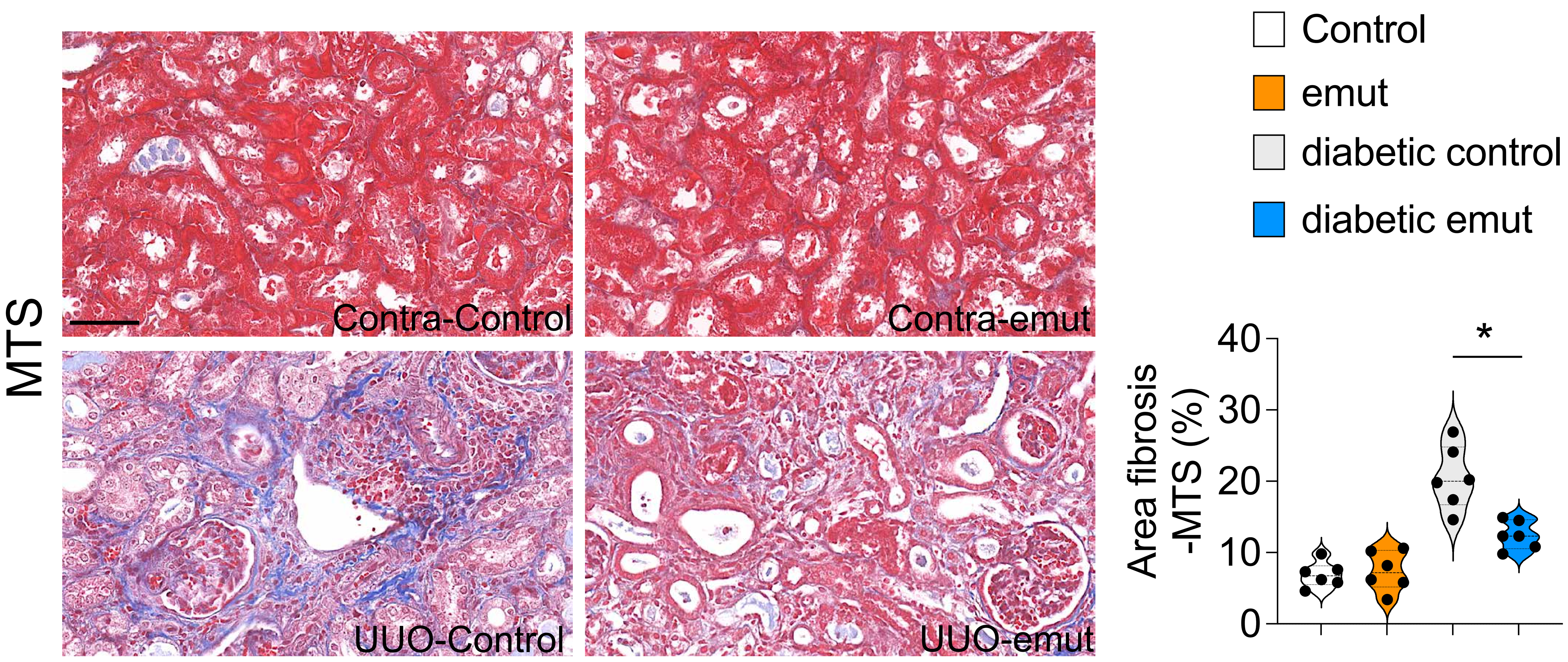
MTS
Contra-Control
Contra-emut
UUO-Control
UUO-emut
Control
emut
diabetic control
diabetic emut
Area fibrosis -MTS (%)
40
30
20
10
0
*

Figure S2

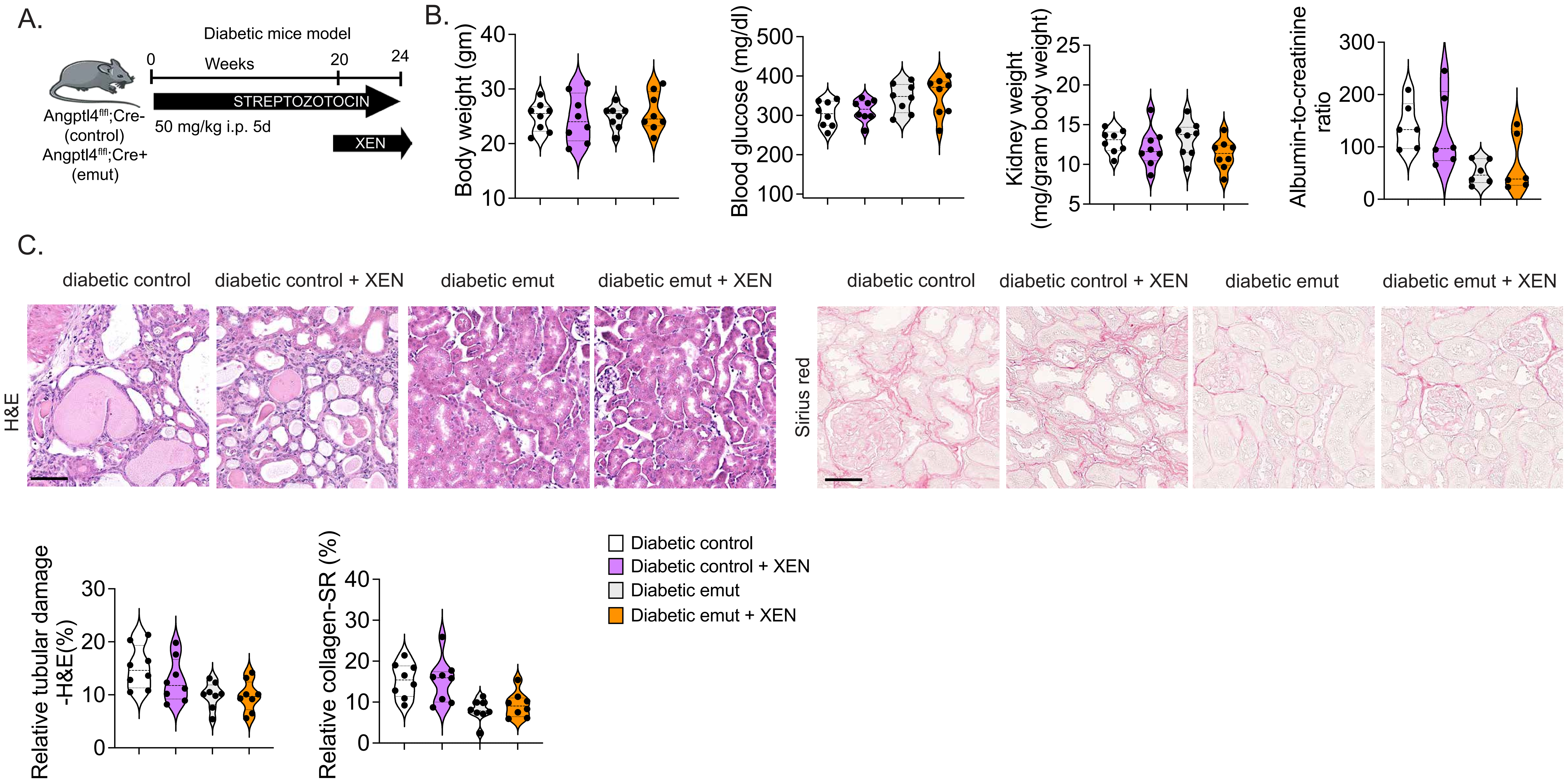
A.
Diabetic mice model
0
Weeks
20
24
STREPTOZOTOCIN
50 mg/kg i.p. 5d
XEN
Angptl4fl/fl;Cre-
(control)
Angptl4fl/fl;Cre+
(emut)
B.
Body weight (gm)
Blood glucose (mg/dl)
Kidney weight
(mg/gram body weight)
Albumin-to-creatinine
ratio
C.
diabetic control
diabetic control + XEN
diabetic emut
diabetic emut + XEN
H&E
Sirius red
Relative tubular damage
-H&E(%)
Relative collagen-SR (%)
Diabetic control
Diabetic control + XEN
Diabetic emut
Diabetic emut + XEN

Figure S3.

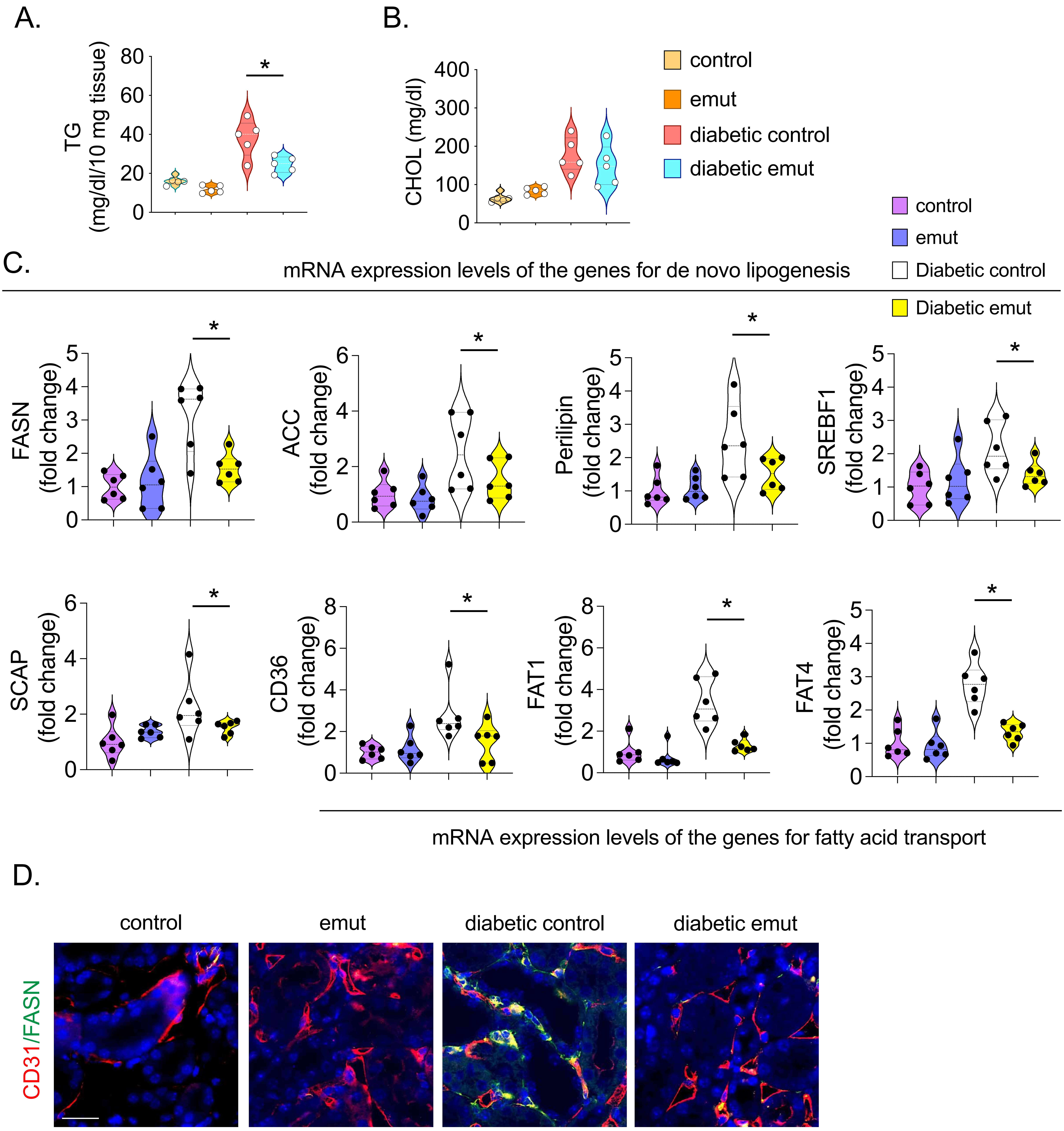
A.
TG (mg/dl/10 mg tissue)
B.
CHOL (mg/dl)
control
emut
diabetic control
diabetic emut
C.
mRNA expression levels of the genes for de novo lipogenesis
control
emut
Diabetic control
Diabetic emut
FASN (fold change)
ACC (fold change)
Perilipin (fold change)
SREBF1 (fold change)
SCAP (fold change)
CD36 (fold change)
FAT1 (fold change)
FAT4 (fold change)
mRNA expression levels of the genes for fatty acid transport
D.
control
emut
diabetic control
diabetic emut
CD31/FASN

Figure S4

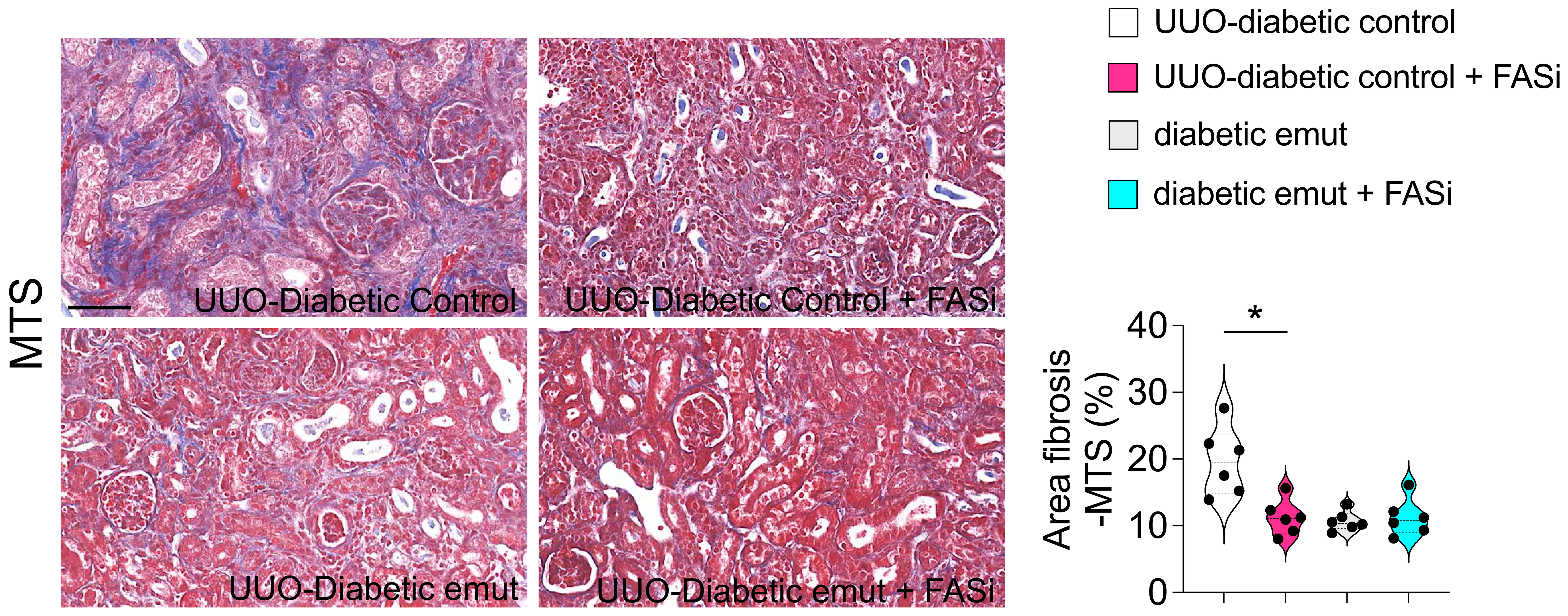

UUO-diabetic control
UUO-diabetic control + FASi
diabetic emut
diabetic emut + FASi
MTS
UUO-Diabetic Control
UUO-Diabetic Control + FASi
UUO-Diabetic emut
UUO-Diabetic emut + FASi
Area fibrosis -MTS (%)
40
30
20
10
0
*


Figure S5

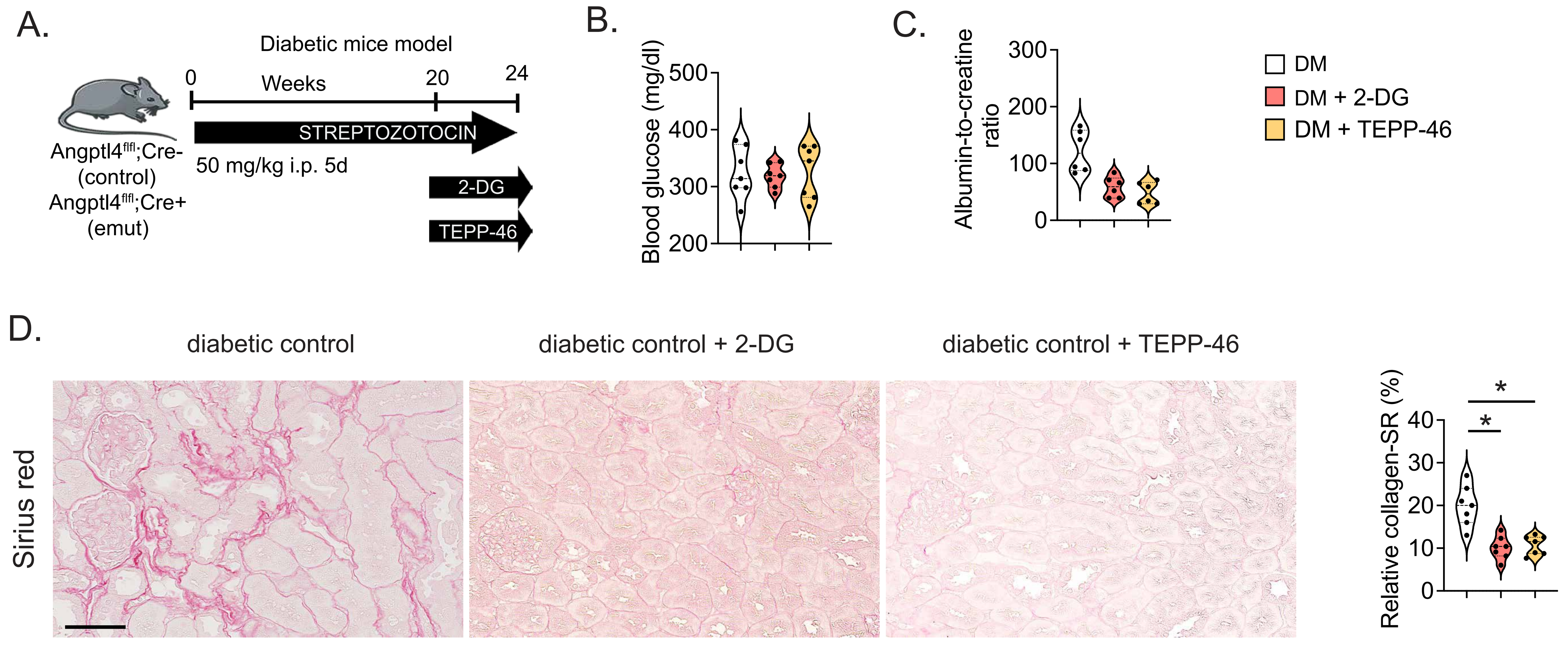
A.
Diabetic mice model
Weeks
0
20
24
STREPTOZOTOCIN
50 mg/kg i.p. 5d
2-DG
TEPP-46
Angptl4fl/fl;Cre-
(control)
Angptl4fl/fl;Cre+
(emut)
B.
Blood glucose (mg/dl)
500
400
300
200
C.
Albumin-to-creatine ratio
300
200
100
0
DM
DM + 2-DG
DM + TEPP-46
D.
diabetic control
diabetic control + 2-DG
diabetic control + TEPP-46
Sirius red
Relative collagen-SR (%)
40
30
20
10
0
*
*

Figure S6.

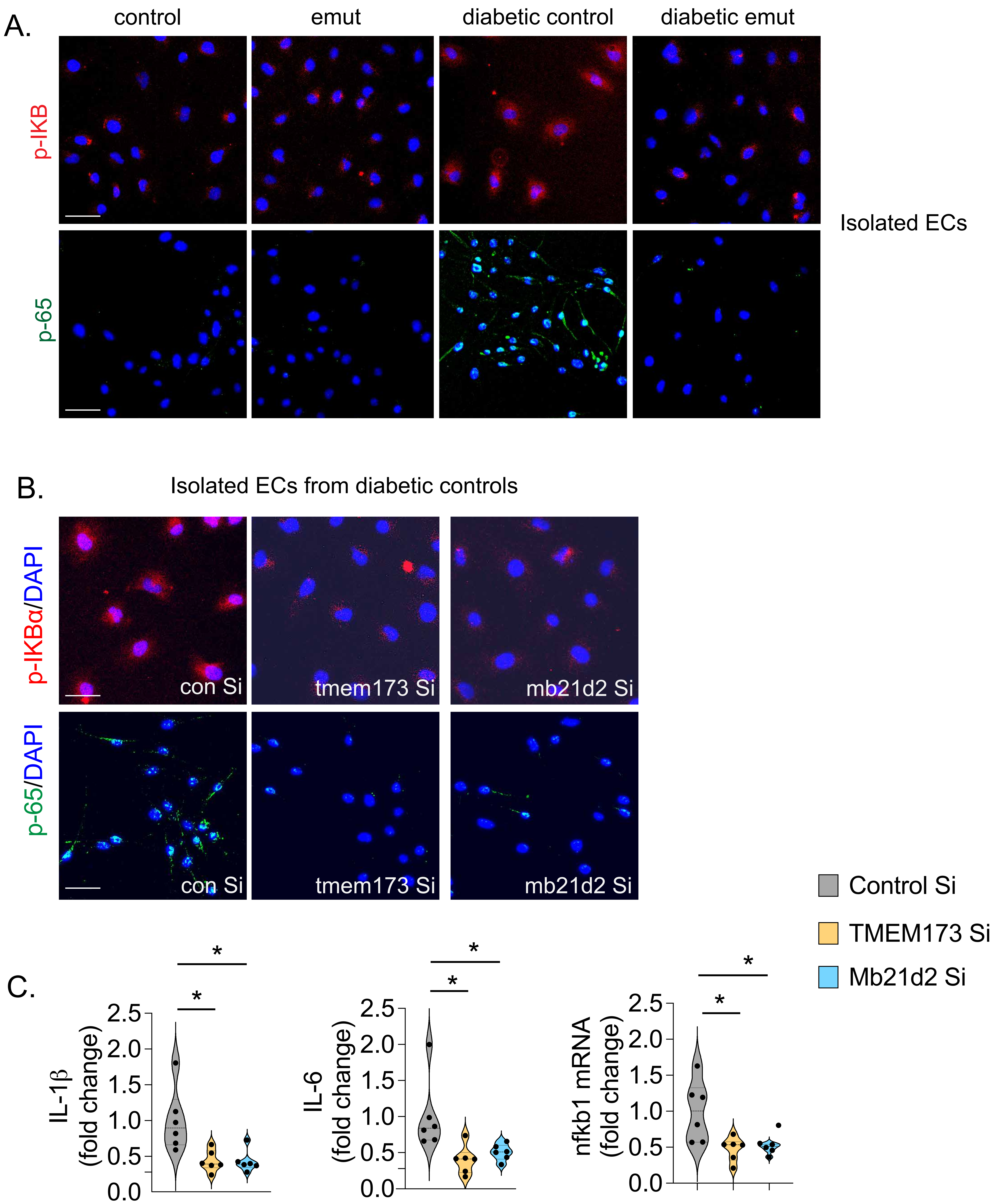
A.
control
emut
diabetic control
diabetic emut
p-IKB
p-65
Isolated ECs
B.
Isolated ECs from diabetic controls
p-IKBα/DAPI
p-65/DAPI
con Si
tmem173 Si
mb21d2 Si
C.
IL-1β
(fold change)
IL-6
(fold change)
nfkb1 mRNA
(fold change)
0.0
0.5
1.0
1.5
2.0
2.5
*
Control Si
TMEM173 Si
Mb21d2 Si

Figure S7

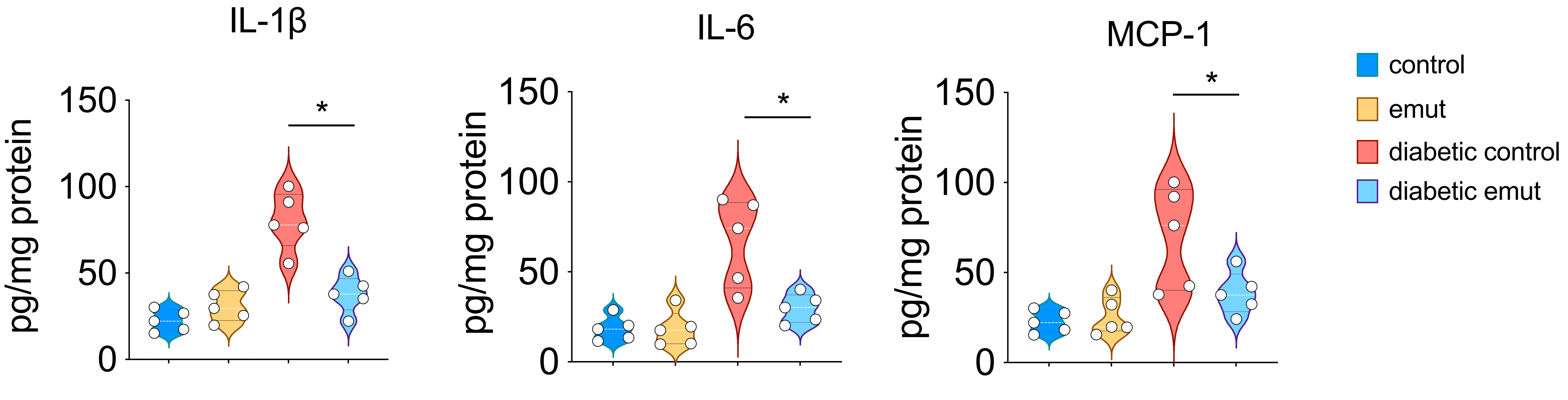
IL-1β
IL-6
MCP-1
pg/mg protein
150
100
50
0
*
control
emut
diabetic control
diabetic emut

Figure S8

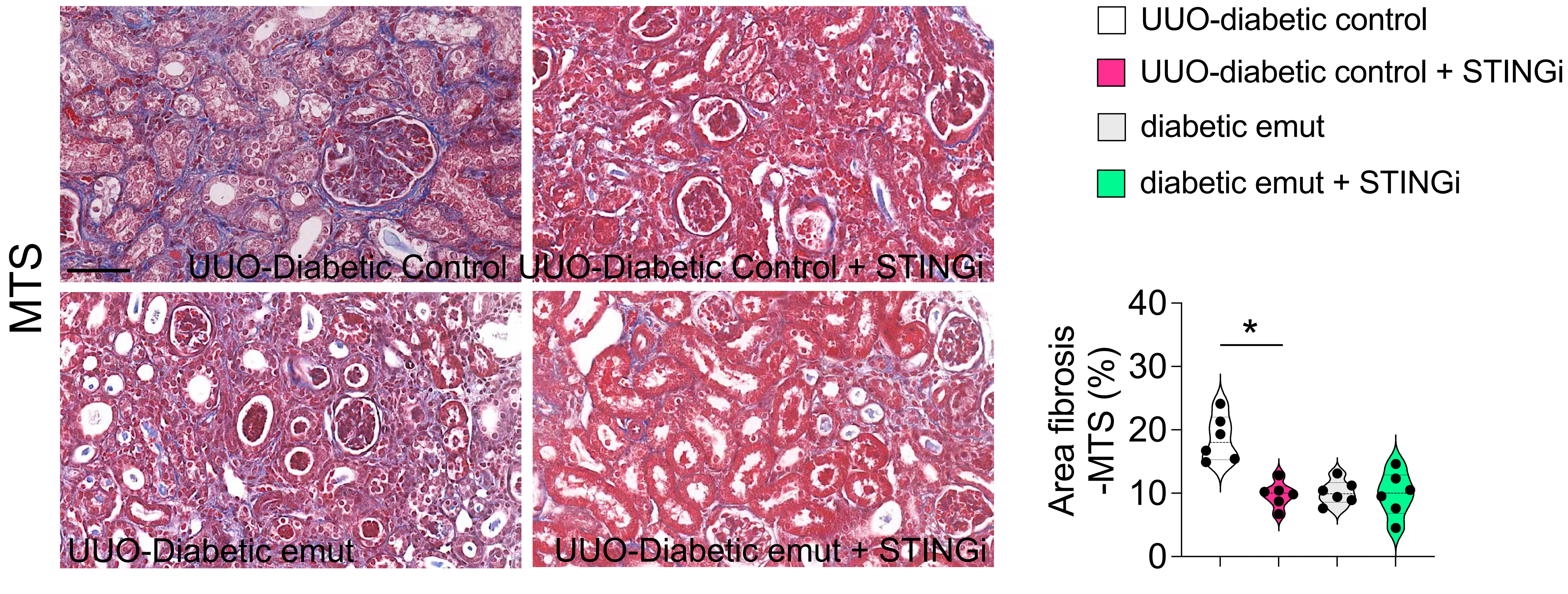
MTS
UUO-Diabetic Control
UUO-Diabetic Control + STINGi
UUO-Diabetic emut
UUO-Diabetic emut + STINGi
UUO-diabetic control
UUO-diabetic control + STINGi
diabetic emut
diabetic emut + STINGi
Area fibrosis -MTS (%)
40
30
20
10
0
*

Figure S9

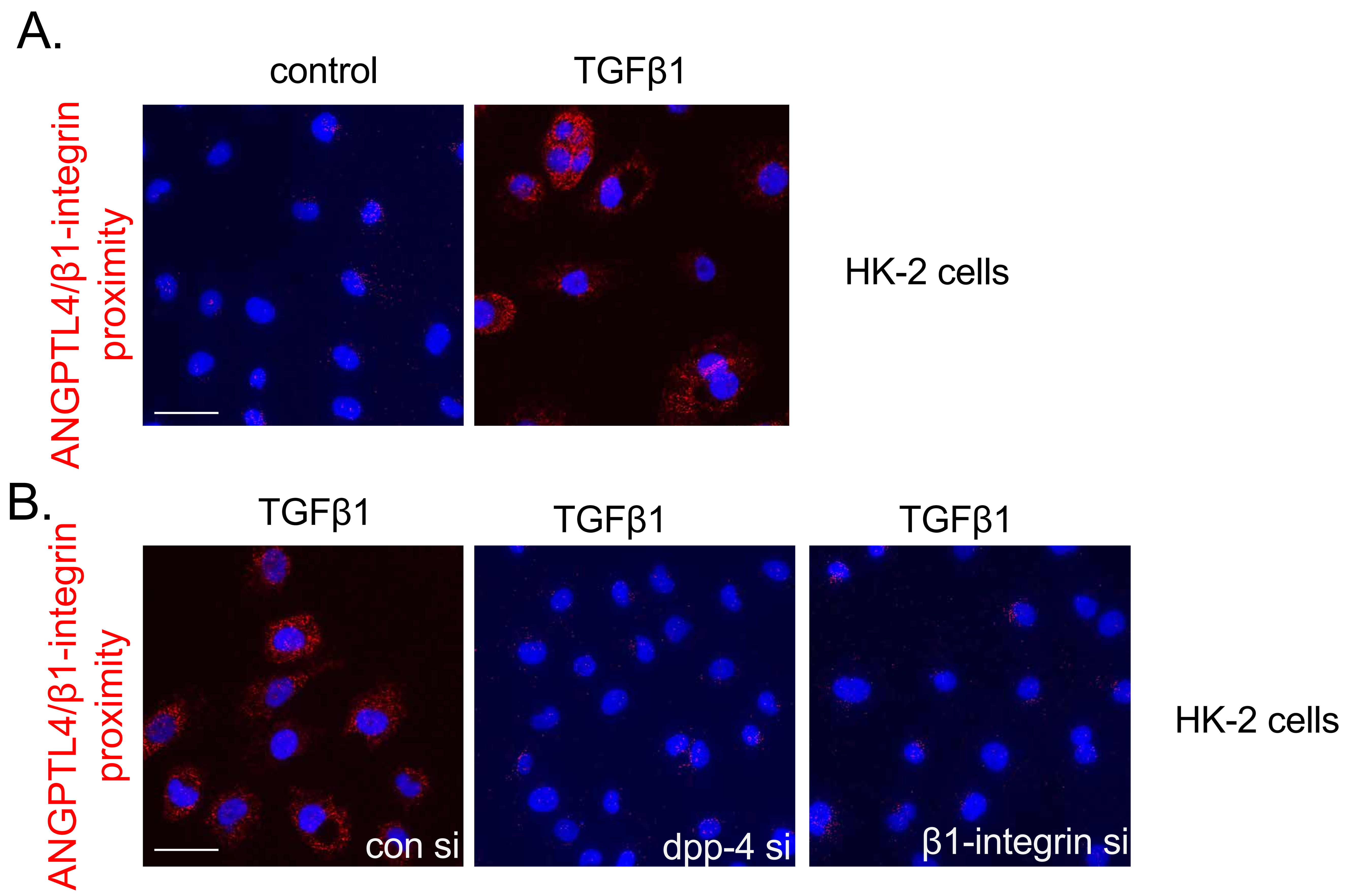
A.
control
TGFβ1
ANGPTL4/β1-integrin proximity
HK-2 cells
B.
TGFβ1
TGFβ1
TGFβ1
ANGPTL4/β1-integrin proximity
con si
dpp-4 si
β1-integrin si
HK-2 cells

Figure S10

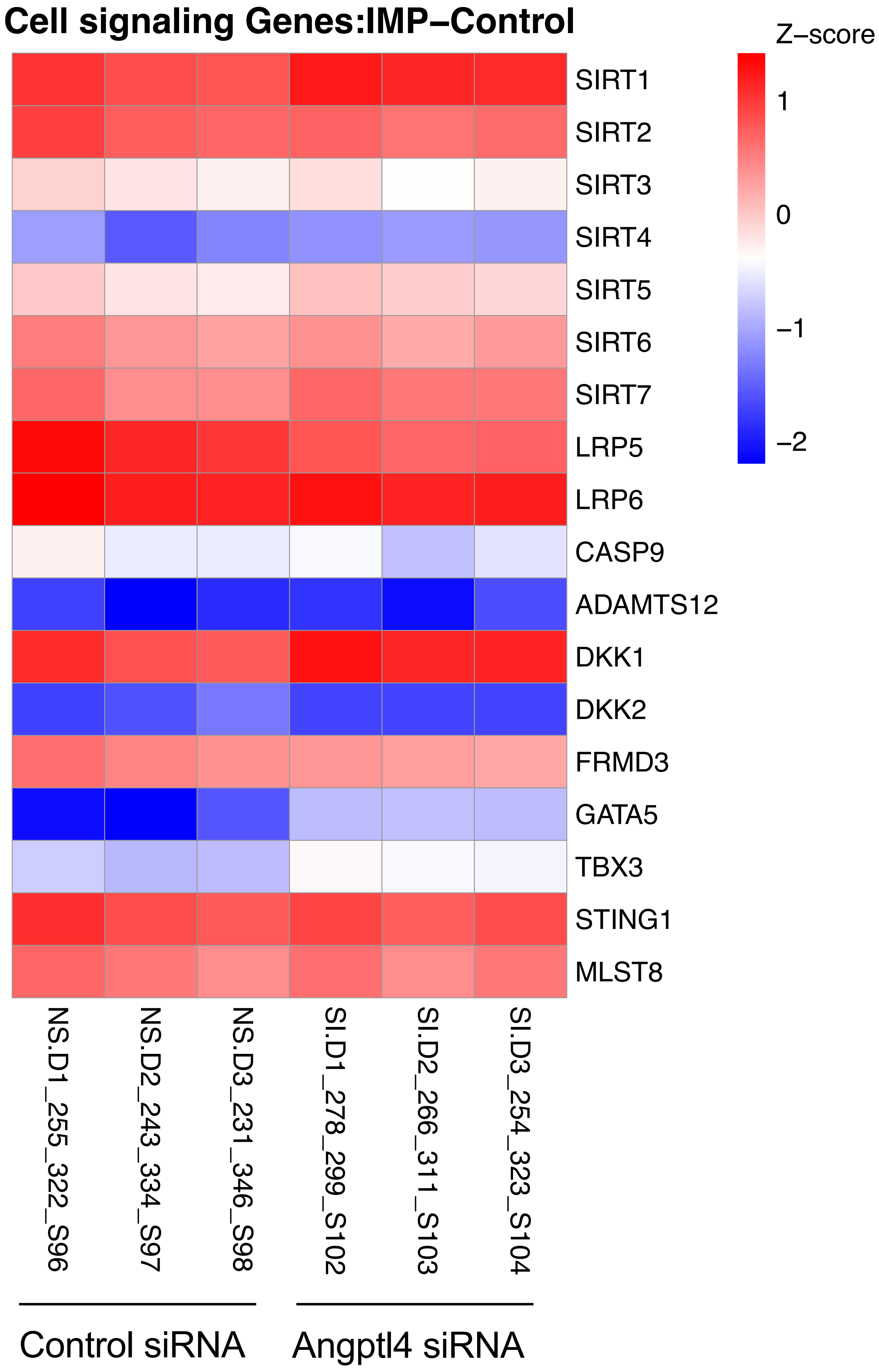
Cell signaling Genes:IMP−Control
Z−score
1
0
−1
−2
SIRT1
SIRT2
SIRT3
SIRT4
SIRT5
SIRT6
SIRT7
LRP5
LRP6
CASP9
ADAMTS12
DKK1
DKK2
FRMD3
GATA5
TBX3
STING1
MLST8
NS.D1_255_322_S96
NS.D2_243_334_S97
NS.D3_231_346_S98
SI.D1_278_299_S102
SI.D2_266_311_S103
SI.D3_254_323_S104
Control siRNA
Angptl4 siRNA

Figure S11

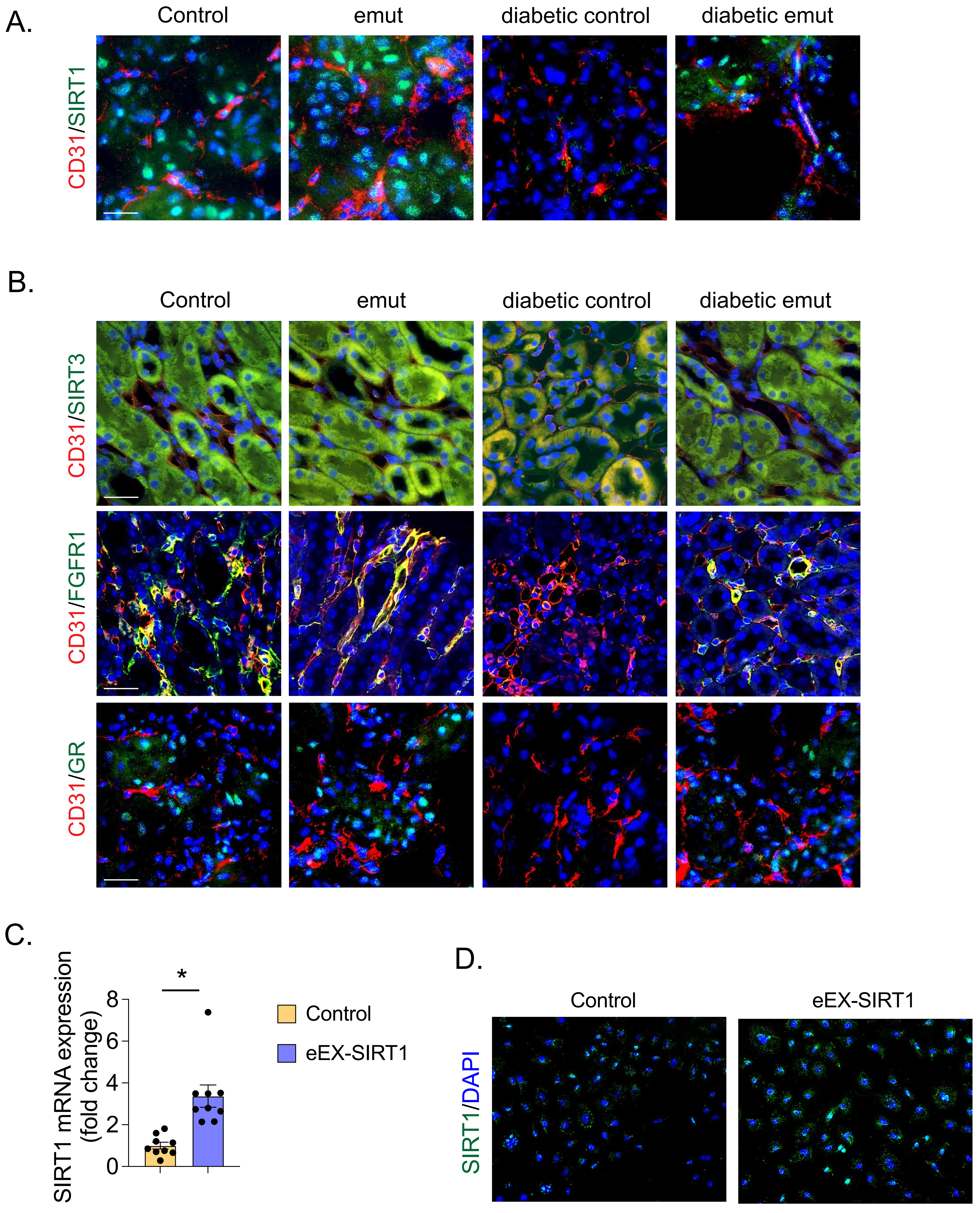
A.
Control
emut
diabetic control
diabetic emut
CD31/SIRT1
B.
Control
emut
diabetic control
diabetic emut
CD31/SIRT3
CD31/FGFR1
CD31/GR
C.
SIRT1 mRNA expression (fold change)
*
8
6
4
2
0
Control
eEX-SIRT1
D.
Control
eEX-SIRT1
SIRT1/DAPI

Figure S12
