## Supplementary material for "Metabolic reprogramming by endothelial ANGPTL4 depletion protects against diabetic kidney disease": Table S1

**Table S1: Sequences mouse primers used**

| Gene name | Forward primer | Reverse primer |
| --- | --- | --- |
| TNFα | 5′-AGGTCCCTACAGGGAACAAA | 5′-TCTCTCTTTCCTTAGACACACGA |
| αSMA | 5′-CTGACAGAGGCACCACTGAA | 5′-GAAATAGCCAAGCTCAG |
| ANGPTL4 | 5’- ACTTCAGATGGAGGCTGGAC | 5’- TCCGAAGCCATCCTTGTAGG |
| ANGPTL3 | 5’- GAGGAGCAGCTAACCAACTTAAT | 5’- TCTGCATGTGCTGTTGACTTAAT |
| ANGPTL8 | 5’- GCTTTACACCTTCGAGCTGA | 5’- ATCCAGGTAGTCTCAGGCTG |
| FSP-1 | 5’-TTCCAGAAGGTGATGAG | 5’-TCATGGCAATGCAGGACAGGAAGA |
| Colla I | 5’-ATCTCCTGGTGCTGATGGAC | 5’- ACCTTGTTTGCCAGGTTCAC |
| IL-6 | 5′-TCTGAAGGACTCTGGCTTTG | 5′-GATGGATGCTACCAAACTGGA |
| TGFβR1 | 5’- CGTGTGCCAAATGAAGAGGAT | 5’- AAGGTGGTGCCCTCTGAAATG |
| Ndufv2 | 5’-TGGATGGCTACCTATCTCCGCT | 5’-GGTACTTCCCAACTGGCTTTCG |
| Cpt1a | 5′- GGTCTTCTCGGGTCGAAAGC | 5′- TCCTCCCACCAGTCACTCAC |
| FASN | 5’- CTTCGCCAACTCTACCATGG | 5’-TTCCACACCCATGAGCGAGT |
| mt-Co1 | 5’- GCCCCAGATATAGCATTCCC | 5’- GTTCATCCTGTTCCTGCTCC |
| mt-Cyb | 5’-AGTAGACAAAGCCACCTTGA | 5’-CCGCGATAATAAATGGTAAG |
| TFAM | 5’- GAGCAGCTAACTCCAAGTCAG | 5’- GAGCCGAATCATCCTTTGCCT |
| TMEM173 | 5’-TTTGCCATGTCACAGGATGC | 5’-ATGAGGCGGCAGTTATTTCG |
| Mb21d1 | 5’-TGGTGGGAAGAGTGGTGATTTC | 5’-TGCATTCCAATGGCAGAAGC |
| 18S | 5’-TTCCGATAACGAACGAGACTCT | 5’-GGCTGAACGCCACTTGTC |

**Table S2: Sequences human primers**

| Gene name | Forward primer | Reverse primer |
| --- | --- | --- |
| ANGPTL4 | 5’- CACAGCCTGCAGACACAACTC | 5’-GGAGGCCAAACTGGCTTTGC |
| ANGPTL3 | 5’- CCTGAAACTCCAGAACACCCAG | 5’- TTCCACGGTCTGGAGAAGGTCT |
| ANGPTL8 | 5’- CAGAAGGTGCTACGGGACAG | 5’- AAATTCTCGGTAGGCAGGGC |
| IL-1β | 5’-CCACAGACCTTCCAGGAGAATG | 5’-GTGCAGTTCAGTGATCGTACAGG |
| IL-6 | 5’-AGACAGCCACTCACCTCTTCAG | 5’-TTCTGCCAGTGCCTCTTTGCTG |
| TNFα | 5’-CTCTTCTGCCTGCTGCACTTTG | 5’-ATGGGCTACAGGCTTGTCACTC |
| 18S | 5’- CAGCCACCCGAGATTGAGCA | 5’- TAGTAGCGACGGGCGGTGTG |
